# Adaptive radiation of barbs of the genus *Labeobarbus* (Cyprinidae) in the East African river

**DOI:** 10.1101/478990

**Authors:** B.A. Levin, M. Casal-López, E. Simonov, Yu.Yu. Dgebuadze, N.S. Mugue, A.V. Tiunov, I. Doadrio, A.S. Golubtsov

## Abstract

Large African barbs of the genus *Labeobarbus* are widely distributed in African freshwaters, and exhibit profound phenotypic plasticity that could be a prerequisite for adaptive radiation. Using morphological, molecular, and stable isotope analyses, we investigated whether an adaptive radiation has occurred in a riverine assemblage of the *L. gananensis* complex. This complex is composed of six phenotypically distinct sympatric forms inhabiting the Genale River (Ethiopian highlands, East Africa). Of the six forms, five were divergent in their mouth morphology, corresponding to ‘generalized’, ‘lipped’, ‘scraping’ (two forms) and ‘large-mouthed’ phenotypes. Stable isotope analysis revealed differences in 15N and 13C among these forms, representing different foraging strategies (omnivorous, scraping and piscivorous). Phylogenetic analysis of two mtDNA markers confirmed the monophyly of *L. gananensis*, suggesting an intra-riverine radiation. However, the Genale assemblage appears to have originated through a combination of allopatric and sympatric events. Some of the specialized forms within this drainage originated independently from the local generalized forms in three different river regions within local ‘mini-flocks’ composed of two to three sympatric forms. Our study shows that adaptive radiation in rivers can be enhanced by a combination of sympatric speciation and temporal geographic isolation, leading to local sympatric speciation followed by migration.

## 1. Introduction

The origin of diversity has intrigued evolutionary biologists for decades. Phenotypic and functional diversity among species or populations can promote resource division and avoidance of inter- and intraspecific competition within a common environment (Schluter 2000). Sympatric adaptive speciation is one of the most important processes that can generate increased diversity and ultimately lead to speciation in the absence of geographic isolation (Kondrashov and Mina 1986; Schliewen et al. 1994; Seehausen and Wagner 2014). Species flocks have repeatedly been reported in lacustrine ecosystems as products of sympatric adaptive radiations (Greenwood 1974; Kontula et al. 2003; Horstkotte and Strecker 2005). For example, there are many outstanding cases of sympatric radiations in cichlid species flocks from the Great Lakes of East Africa (Meyer et al. 1990; Salzburger et al. 2002; Seehausen et al. 2003) and from small crater lakes (Schliewen et al. 1994; Barluenga et al. 2006; Franchini et el. 2014). In all of these cases, resource partitioning was found to be the main factor driving the divergence. In resource-poor habitats such as postglacial lakes and crater lakes, speciation typically occurs through disruptive ecological selection (Schluter 2000; Elmer et al. 2010). Some of the well-known cases of ecological divergence in lake isolates include the benthic and limnetic forms of three-spined stickleback (Schluter 1996; McKinnon and Rundle 2002), whitefish (Østbye et al. 2006; Laporte et al. 2015), and the various forms of Arctic charr (Alekseyev et al. 2002; Gordeeva et al. 2014).

There are also striking examples of ecological divergence in sympatry among cyprinids, the most species-rich teleostean family. A presumed species flock of small Asian ‘*Barbus*’ from Lake Lanao in the Philippines was composed of 18 to 20 species (Herre 1933; Myers 1960); all but one became extinct due to human activities before genetic tools had been developed (Kornfield and Echelle 1984). Similarly, the Altai Osman of the genus *Oreoleuciscus* from the Valley of the Lakes in Mongolia has diverged into sympatric dwarf and piscivorous forms (Dgebuadze 1995). Finally, an outstanding cyprinid species flock of large African barbs (genus *Labeobarbus*) was described from Lake Tana, Ethiopia, which is comprised of 15 phenotypically and ecologically divergent sympatric forms or species (Nagelkerke et al. 1994; Mina et al. 1996; de Graaf et al. 2008).

While sympatric speciation has been studied extensively in lacustrine systems, less research has been conducted in riverine environments (*i.e*. more open to gene flow; Castric et al. 2001). Nevertheless, some remarkable examples of riverine radiations have recently been discovered in different fish groups, such as mormyrids (Sullivan et al. 2002; Feulner et al. 2007), cyprinids (Roberts 1998; Dimmick et al. 2001), and cichlids (Koblmüller et al. 2008; Schwarzer et al. 2011; Piálek et al. 2012, 2018). The formation of species flocks among distant lineages may indicate independent origins of the various species, and suggests that some conditions in riverine environments may promote sympatric speciation.

A putative riverine species flock was discovered over twenty years ago in the Ethiopian cyprinid genus *Labeobarbus* in the Genale River of Ethiopia (Golubtsov 1993; Mina et al. 1998). This riverine assemblage of the Genale barbs was comprised of fewer distinct forms than those found in the lacustrine species flock from Lake Tana (Golubtsov 1993). Divergent morphology and preliminary gut content analysis of these forms suggested trophic specialization (Mina et al. 1998; Golubtsov 2010). Specifically, five different forms were reported, which included generalized, thick-lipped, large-mouthed (piscivorous), as well as two scraping forms with different jaw scrapers (Golubtsov 1993; Mina et al. 1998). These same five forms were recently sampled again, along with an additional new form named ‘short’ due to its short body (Fig. 1). Preliminary studies have documented genetic divergence among some of these sympatric Genale barb forms (Dimmick et al. 2001; Levin et al. 2013). This is in contrast to the well-studied *Labeobarbus* species flock in Lake Tana, which displays no or shallow genetic divergence among sympatric forms (de Graaf et al. 2010; Beshera and Harris 2014; Nagelkerke et al. 2015). The assemblage of large African barbs in the Genale River consists of species from two different genera: *Labeobarbus* (*L. gananensis*) and *Varicorhinus* (*V. jubae*). However, it was recently reported that the scraper form *V. jubae* and generalaized *L. gananensis* are sister species, suggesting that the scraping foraging strategy has evolved independently among barbs, and that the previously assigned ‘*Varicorhinus’ jubae* belongs to the genus *Labeobarbus* (Levin et al. 2013). This finding raises the question as to whether the observed diversity of *Labeobarbus* in the Genale River represents a species flock (*i.e*. the species are monophyletic), or a species stock (*i.e*. an assemblage of allopatrically originated forms).

**Fig. 1a.**
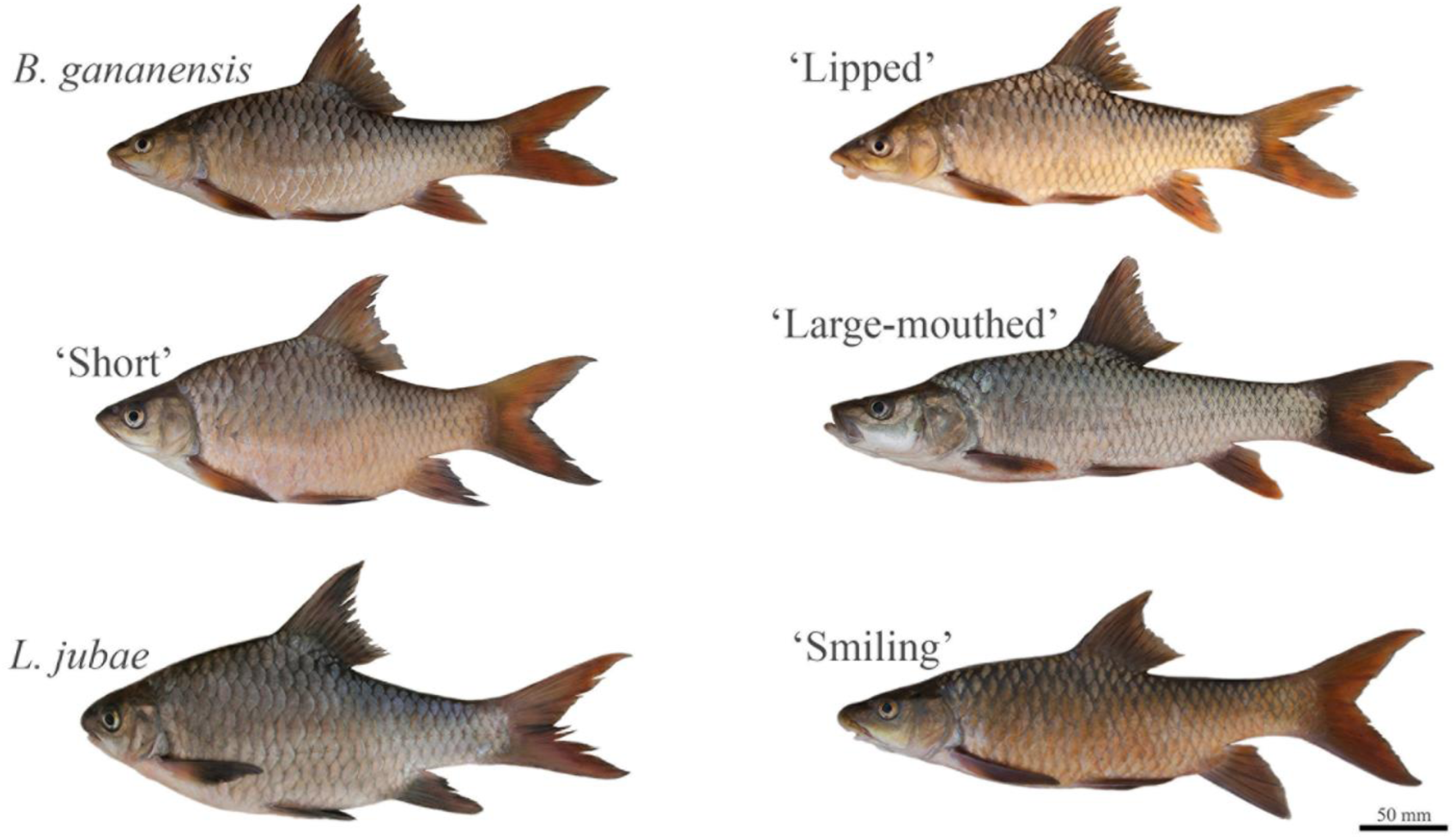
Lateral view of the *Labeobarbus* forms from the Genale River.

**Fig. 1b.**
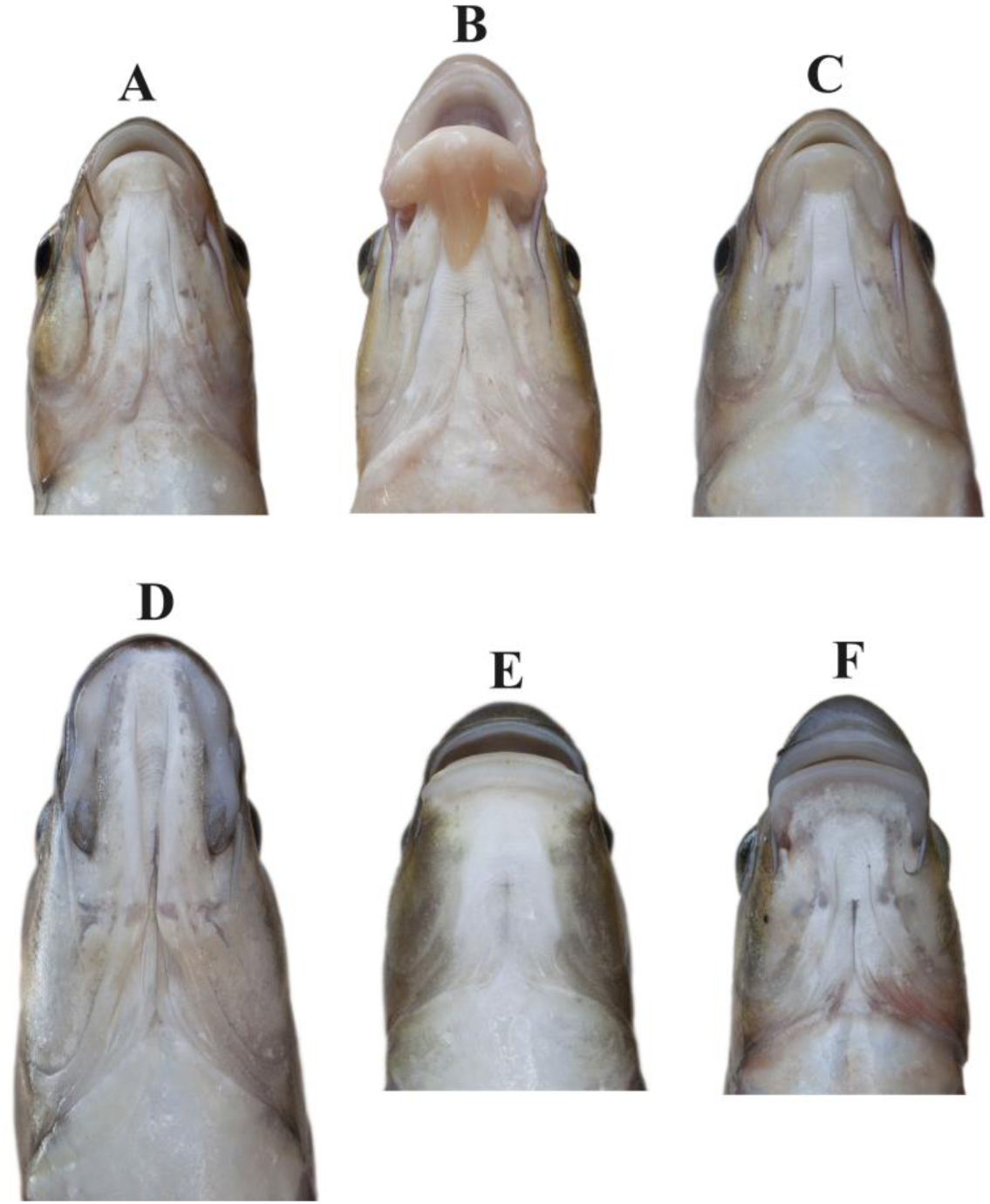
Mouth features of Genale barbs: A – ‘Generalized’ form of *L. gananensis*, B – ‘Lipped’ form of *L. gananensis*, C – ‘Short’ form of *L. gananensis*, D – ‘Large-mouthed’ form (*L.* sp.1; piscivorous), E – *L. jubae* (scraper), and F – ‘Smiling’ form (*L.* sp.2; scraper).

The main questions we addressed in this study were: (i) are the phenotypic features found in Genale barbs correlated with the partitioning of trophic resources, and (ii) what is the origin of the observed diversification among Genale barbs? In order to answer these questions, we tested the hypothesis of a monophyletic origin of the Genale barbs as a species flock following sympatric speciation. An alternative hypothesis would be that sympatric riverine barbs are a stock (assemblage) of different lineages that originated in isolated or semi-isolated parts of different river basins (allopatric model of speciation), and later colonized and established self-reproducing populations in sympatry with other Genale river forms. Hence, our specific objectives were to: 1) test the correlation between phenotypic divergence and trophic traits among riverine Genale forms; 2) assess trophic resource partitioning using stable isotope analyses; 3) evaluate the genetic relationships among Genale barbs and samples from other tributaries of the Juba, Wabe-Shebelle, and adjacent basins using mitochondrial markers.

## 2. Materials and methods

### 2.1. Sampling

Fishes of a putative species flock were sampled in the Genale River (Fig. 2), Ethiopia, during March-April 2009 in the framework of the Joint Ethio-Russian Biological Expedition (JERBE). Names of the different forms are described in Table. Comparative samples were collected from six other locations in the Juba and Wabe-Shebelle (JWS) basins (Fig. 2) during 2009-2012, with sampling permission from the appropriate authorities. Fish were caught using cast and gill nets, and killed with an overdose of MS-222. Samples were first preserved in 10% formalin in the field, and then transferred to 70% ethanol. All specimens (Supplementary Table S1) are stored in the Zoological Museum of Moscow University (ZMMU).

**Table.** Common names of the six forms of African barbs from the Genale River, and the preliminary qualitative descriptions used in the field to identify each form.

| Latin name with our specification | Name used in the text | Description based on general appearance |
| --- | --- | --- |
| *Labeobarbus gananensis*, 'generalized' form | 'NEC' | Absence of some specialized characters compared to other forms. NEC – no extreme characters (Nagelkerke et al. 1994; Mina et al. 1998). Moderately developed, variable lips that diminish towards the middle lobe of the lower lip. |
| *L. gananensis*, 'lipped' form | 'Lipped' | Greatly developed lips, referred to as 'rubber lips' (Matthes 1963), 'thick-lipped' or 'hypertrophic' in cichlids (Machado-Schiaffino et al. 2014). Intermediate lobe of lower lip is well developed and unattached. |
| *L. gananensis*, 'short' form | 'Short' | Short and deep-bodied, rarely possessing enlarged lips. |
| *L. jubae* | 'Jubae' | Scraping form with short head and relatively deep body. Mouth is inferior and wide. Lower jaw modified |
|  |  | into scraper, covered by horny cutting sheath. Lower lip is drastically reduced. Barbels are short. |
| *L. sp.1*, scraping form<br>'smiling' | 'Smiling' | Scraping form with slender body. Mouth is inferior and wide. Lower jaw modified into scraper different than that of *L. jubae*. Snout is enlarged with well-expressed rostral cap. |
| *L. sp.2*, piscivorous form | 'Large-mouthed' | Slender body, elongated head with terminal mouth and significantly elongated jaws. Most specimens have a weakly expressed arch behind the head. Middle lobe of lower lips is absent. |

**Fig. 2.**
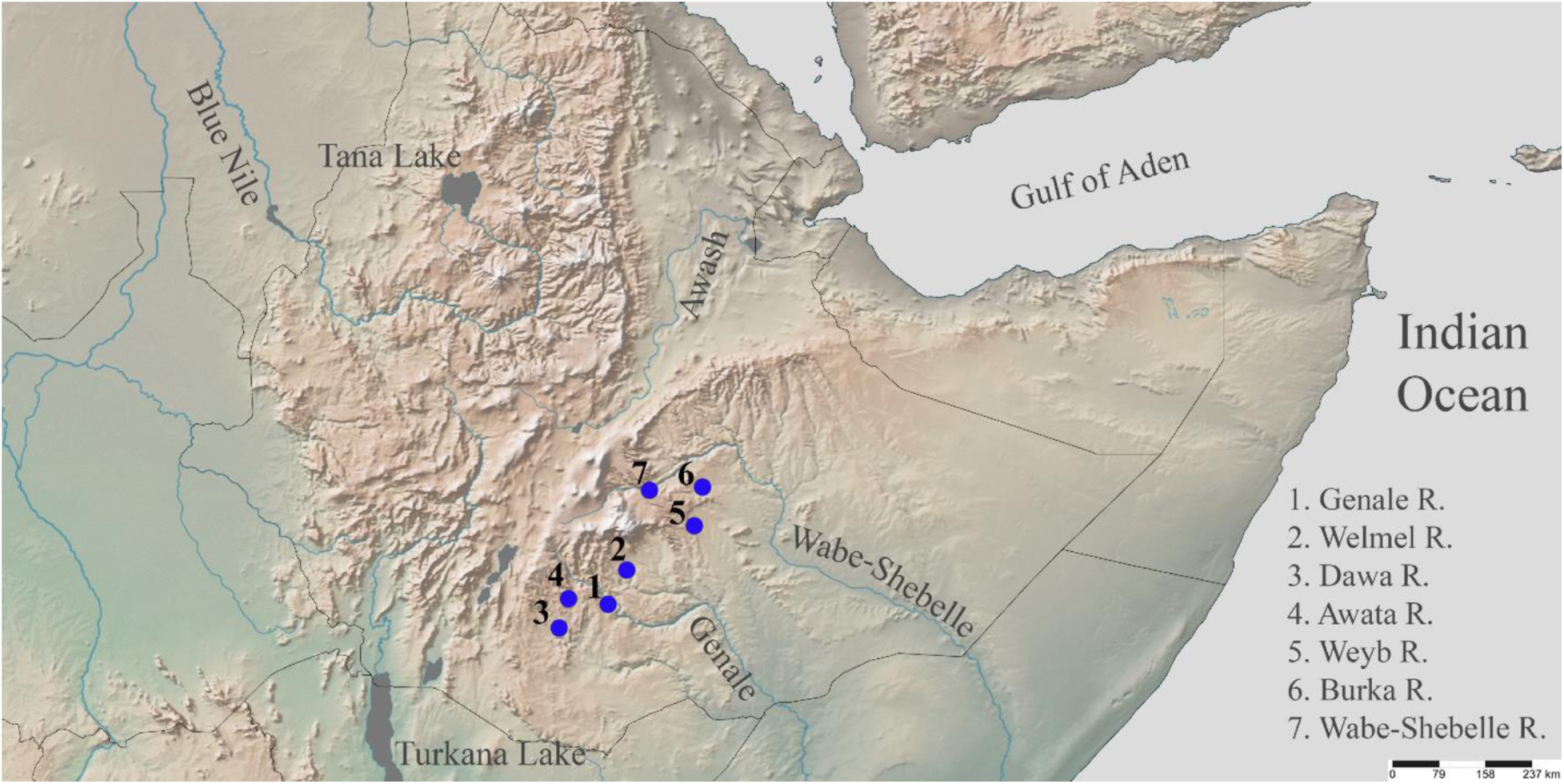
A map of the sampling localities.

### 2.2 Morphological analysis

The following morphometric characters were examined: standard length (SL) and body height (H), measured with a digital caliper (to nearest 0.01 mm), and gut length (GL), measured with a ruler (to nearest 1 mm). Gill arches were stained with Alizarin red, and gill rakers (GR) on both the lower and upper arches were counted using a binocular microscope (Leica EZ4D). Sample size for different traits varied from 97 (GR) to 151 (GL).

### 2.3. Stable isotope analysis

In total, 116 barb samples were processed for isotope analysis. Other species of fishes specialized for various diets were used as references. These included detritivorous cyprinids *Labeo* cf. *cylindricus* (n=12), bentophagous mormyrids *Mormyrus caschive* (n=9), and predatory catfish *Bagrus urostigma* (n=3). Muscle tissue from the dorsal side of the body under the dorsal fin was sampled from freshly collected specimens. Samples were dried at 60ºC and wrapped in tin capsules. Stable isotope analysis was conducted at the Joint Usage Center of the A.N. Severtsov Institute of Ecology and Evolution RAS, Moscow. Briefly, a Thermo Delta V Plus continuous-flow IRMS was coupled with an elemental analyzer (Flash 1112) and equipped with a Thermo No-Blank device. The isotopic composition of N and C was expressed in a δ-notation relative to the international standards (atmospheric nitrogen and VPDB, respectively): δX (‰) = [(R_sample_/R_standard_)-1] × 1000, where R is the ratio of the heavier isotope to the lighter. Samples were analyzed with reference gas calibrated against the International Atomic Energy Agency (IAEA) reference materials USGS 40 and USGS 41 (glutamic acid). Measurement accuracy was ± 0.2 δ units. Along with isotopic analysis, nitrogen and carbon content (as %) and C/N ratios were determined in all samples. The C/N (mass/mass) differed slightly among the different forms, with an average of 3.20 (SD = 0.09). This indicates that the lipid concentration was uniformly low, and no lipid normalization of δ^13^С values was required (Post et al. 2007).

### 2.4. Morphology and stable isotope statistics

A non-parametric Spearman's rank correlation coefficient was used for testing the association between variables. Dunn’s *post hoc* test was used following a non-parametric Kruskal-Wallis test, and results were visualized with boxplots constructed using R packages (v. 3.3.2). Analysis of variance (ANOVA) and Tukey’s unequal n honestly significant difference (HSD) were used to compare δ^13^С and δ^15^N values in the different forms. All statistical analyses were performed using Statistica 8.0.

### 2.5. DNA sampling, extraction, amplification, and sequencing

A total of 159 Genale barbs were used for DNA analyses, including all forms of the *L. gananensis* complex and *L. jubae* from the Genale watershed. An additional 40 DNA samples of *Labeobarbus* were included from six other locations in Indian Ocean catchment – four from different tributaries of the Juba basin, and two from the Wabe-Shebelle basin (Supplementary Table S1, Fig. 2). Sequences from an additional five species of *Labeobarbus*, as well as the closely-related specialized scraper *Varicorhinus beso*, were retrieved from GenBank for comparative purposes. These specimens originated from adjacent drainages of the Atlantic (Tana Lake and Blue Nile) and Indian Ocean (Tana River basin and Malawi Lake) catchments, and inland basins (Rift Valley lakes and Omo-Turkana Basin) (Supplementary Table S2). *Carasobarbus fritschii* and *Enteromius ablabes* were included as outgroups.

Total genomic DNA was extracted from ethanol-preserved fin tissues using the BioSprint 15 kit for tissue and blood (Qiagen). A 1049 bp fragment of the mtDNA cytochrome *b* gene (cyt-*b*) and 615 bp fragment of the control region (*d-loop*) were amplified (Supplementary Table S3; Meyer et al. 1994; Palumbi 1996; Doadrio and Perdices 2001; Levin et al. 2013). PCR products were visualized on 1% agarose gels, purified with ExoSAP-IT™ and sequenced by MACROGEN (Seoul, Republic of Korea) using ABI 3730XL, or at the Institute of Biology of Inland Waters (Russian Academy of Sciences) using ABI 3500. All new sequences were deposited in GenBank (Accession Numbers: MK001025-MK001392 and MK015650-MK015652, see Supplementary Table S1).

### 2.6. Sequence analysis

All sequences were aligned and edited using Clustal X (Thompson et al. 1994) as implemented in MEGA v. 6.0 (Tamura et al. 2013). In total, 191 specimens from seven rivers in the JWS riverine systems were analysed for cyt-*b*; 222 were analysed for *d-loop*.

### 2.7. Phylogenetic analysis and haplotype network construction

All sequences were collapsed into common haplotypes using ALTER software (Glez-Peña et al. 2010). Both mtDNA genes were concatenated for construction of a multi-locus phylogeny. The best-fit model of molecular evolution by codon position was estimated via Akaike information criterion (AIC; Akaike 1973) using PartitionFinder v. 1.1.1 (Lanfear et al. 2012). The best partition schemes used in the different phylogenetic analyses are presented in Supplementary Table S4.

Bayesian phylogenetic inference (BI) was carried out in MrBayes v. 3.2.6 (Ronquist et al. 2012). Two simultaneous analyses were run for 1 600 000 generations (set 1; cyt-*b*) and 4 000 000 generations (set 2; concatenated cyt-*b* + *d-loop*), each with four MCMC chains sampled every 100 generations. Convergence was verified with Tracer v. 1.6. (Rambaut et al. 2014). A 50% majority rule consensus tree with posterior probabilities was obtained after the first 25% of generations were discarded as burn-in. Phylogenetic analyses of both datasets was also conducted with maximum likelihood (ML) in RaxML (Stamatakis 2007). The GTRGAMMAI substitution model was applied with 1000 replicates using the rapid bootstrap algorithm (Stamatakis et al. 2008). Uncorrected *p*-distances for both genes were calculated in MEGA v. 6.0 (Tamura et al. 2013) in order to quantify genetic divergence among forms and populations.

Bayes factor (BF) comparisons were used to test for monophyly of the JWS lineage. The stepping-stone sampling algorithm implemented in MrBayes was used to estimate marginal likelihoods for two alternative models where JWS barbs were constrained to be either monophyletic or paraphyletic. BF was calculated in log-units as the difference between natural logarithms of marginal likelihoods of the two models. The same approach was used to test for monophyly/polyphyly of the ‘NEC’ form of Genale barbs. In both cases, the constrained stepping-stone analysis involved 50 steps and the same number of chains and generations as the corresponding Bayesian phylogenetic analysis (see above), with the initial burn-in of one step.

A haplotype network for the *d-loop* dataset was constructed using the median joining algorithm (Bandelt et al. 1999) in PopArt 1.7 (Leigh and Bryant 2015).

### 2.8. Genetic diversity, population structure and demography

Nucleotide and haplotype diversity parameters were estimated using DnaSP v. 5.0 (Librado and Rozas 2009). Genetic differentiation among populations was tested for in Arlequin v. 3.5.1.2. (Excoffier and Lischer 2010) using the fixation index *F*_ST_ (Weir and Cockerham 1984). To detect signatures of demographic shifts in recent history, deviations from a model of mutation-drift equilibrium were tested for in both genes, using Fu’s *F*_*s*_ (Fu 1997) with 1000 pseudo-replications, as well as Tajima’s *D* neutrality test (Tajima 1989). Both tests were performed in Arlequin v. 3.5.1.2. (Excoffier and Lischer 2010). The critical values obtained by simulations in these neutrality tests were used to assess statistical significance (*p* < 0.05); however the *p*-value was adjusted to 0.02 for Fu’s *Fs* due to the non-normal distribution of the *Fs* statistic (Ramos-Onsins and Rozas 2002).

## 3. Results

### 3.1. Morphology associated with trophic resource partitioning

#### 3.1.1. Gill rakers

Most of the Genale forms were divergent in the number and shape of gill rakers (Fig. 3; Supplementary Fig. S1 and Table S5). GR number did not increase with SL in any form (*r*_*s*_ varied from −0.281 for ‘NEC’ to 0.312 for ‘smiling’, *p* > 0.05). The ‘large-mouthed’ form had significantly fewer GR compared to most of the other forms (Kruskall-Wallis test, *p* < 0.05), except the ‘lipped’ form (*p* = 0.18). Both scraping forms (*L. jubae* and ‘smiling’) had higher GR numbers compared to the other forms (Kruskall-Wallis test, *p* < 0.05); the number of GR was similar between the two scraping forms. The ‘NEC’, ‘lipped’ and ‘short’ forms possessed an intermediate and similar number of GR. With the exception of the ‘large-mouthed’ form, GR were saturated with taste buds located on processes, serrations or papillae on the pharyngeal side; they were particularly well developed in both scraping forms (Supplementary Fig. S1).

**Fig. 3.**
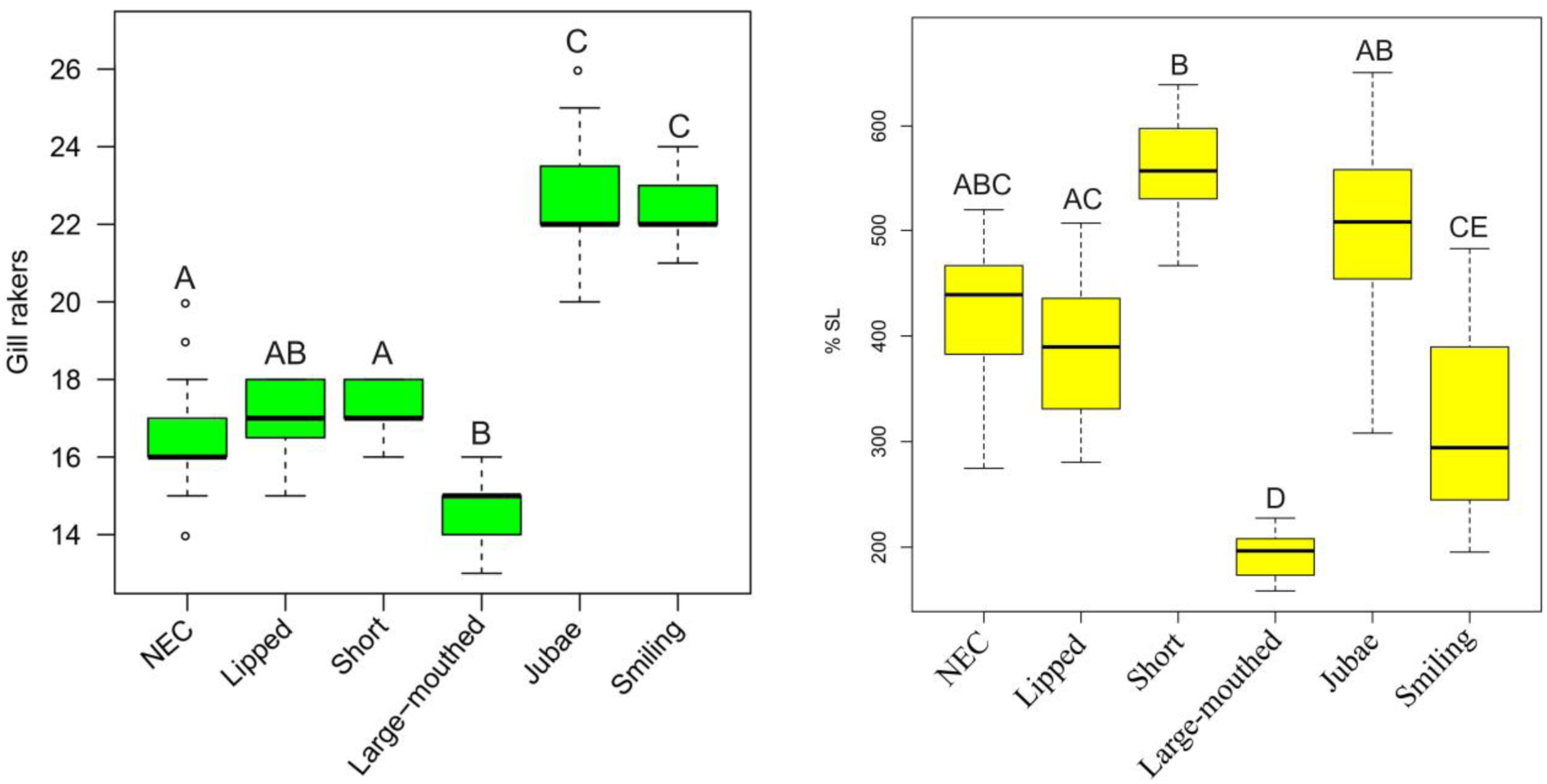
Distribution of gill raker numbers (left) and gut length (right) among Genale forms. Median is shown as the horizontal black line inside the box. The box represents 1^st^ and 3^rd^ quartiles of variation. Different letters above the whiskers indicate significant differences between the forms (*p* < 0.05, Kruskal-Wallis test)

#### 3.1.2. Gut length

With the exception of the ‘large-mouthed’ form, all other forms had long guts that measured from four to six times the body length (Fig. 3). Gut length (GL as % SL) in all forms was positively correlated with body length, with the exception of the ‘large-mouthed’ form that showed a negative (non-significant) correlation (Table S6). The ‘Large-mouthed’ form differed significantly in GL from all other forms; there were significant differences between some of the other forms (Kruskall-Wallis test, *p* < 0.05; Fig. 3). The ‘short’ form appeared to have the longest gut, however the absolute length was similar to the other forms; when expressed in relation to SL, the gut appeared longer because body length was shorter (Fig. 3).

### 3.2. Diet analyses

Preliminary inspection of gut contents revealed that all forms, with the exception of the ‘large-mouthed’, always had full guts, primarily with rich muddy material (detritus with chitinized insect and plant remnants). Several individuals of the ‘smiling’ form had semi-filled guts, containing larvae of benthic Simulidae. Individuals of the ‘large-mouthed’ form had predominantly empty guts; when they were filled, they contained remnants of cyprinid juveniles and adult mochokid catfish *Chiloglanis* sp.

Stable isotope analysis confirmed a considerable difference in trophic niches between certain forms. δ^13^С values were similar in all forms (ANOVA: *p* > 0.3), but significant differences were found in δ^15^N values (F_5,103_ = 69.4, *p* < 0.0001). ‘NEC’, ‘lipped’, and ‘short’ forms had the lowest mean δ^15^N values (< 11.5‰; Fig. 4). The ‘large-mouthed’ form was the most enriched in ^15^N (δ^15^N = 13.8±0.1‰), indicating that it occupied the highest trophic level (Fig. 4). Both scraping forms (‘smiling’ and *L. jubae*) had significantly lower δ^15^N values than the ‘large-mouthed’ form (12.7±0.1‰ and 12.0±0.1‰), but were significantly enriched in ^15^N relative to the ‘NEC’, ‘lipped’, and ‘short’ forms (Fig. 4). Among the scraping forms, ^15^N was higher in the ‘smiling’ form than *L. jubae* (Fig. 4).

**Fig. 4.**
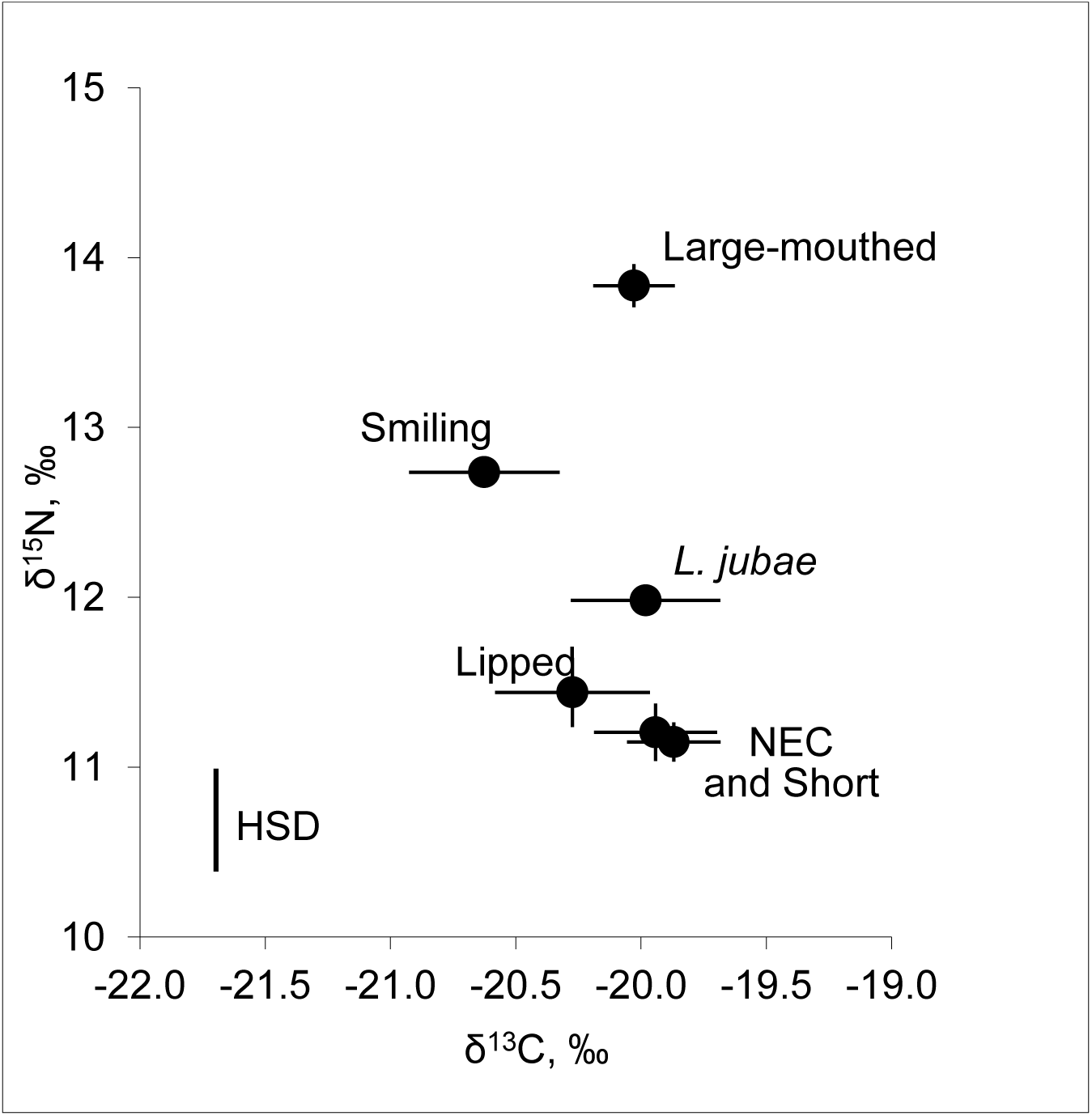
Stable isotope composition (mean δ^13^С and δ^15^N values [±SE], n = 14 - 26) of the six Genale barb forms. HSD: Tukey’s unequal n honestly significant difference for δ^15^N values.

### 3.3. Testing the monophyly of African barbs from the Juba and Wabe-Shebelle (JWS) basins using cyt-b

56 unique haplotypes were found from the JWS basins, which formed a monophyletic clade in both the BI and ML phylogenetic trees (Fig. 5). Monophyly of this group was strongly supported by the topology hypothesis testing (log BF = 13.2). This clade formed a sister clade to the Ethiopian barbs from the Ethiopian Rift Valley and other westward drainages, including those from Lake Tana that we assigned as *L. intermedius* s. lato following Banister (1973). However, the statistical support for this relationship was weak. One specimen from the Arer River (Wabe-Shebelle drainage; GenBank sequence GQ853247) grouped with the western barbs *L. intermedius* s. lato. This could simply be a result of geographic mislabeling of the sample, especially in light of the fact that there is a Hare River (Lake Chamo basin), which could be acoustically mistaken for Arer River (Wabe-Shebelle basin). Alternatively, it could be a result of the penetration of *L. intermedius* into the upper streams of the Wabe-Shebelle drainage by riverine capture(s). It seems more likely that the positioning of the GQ853247 sample in the phylogeny is a labeling error.

**Fig. 5.**
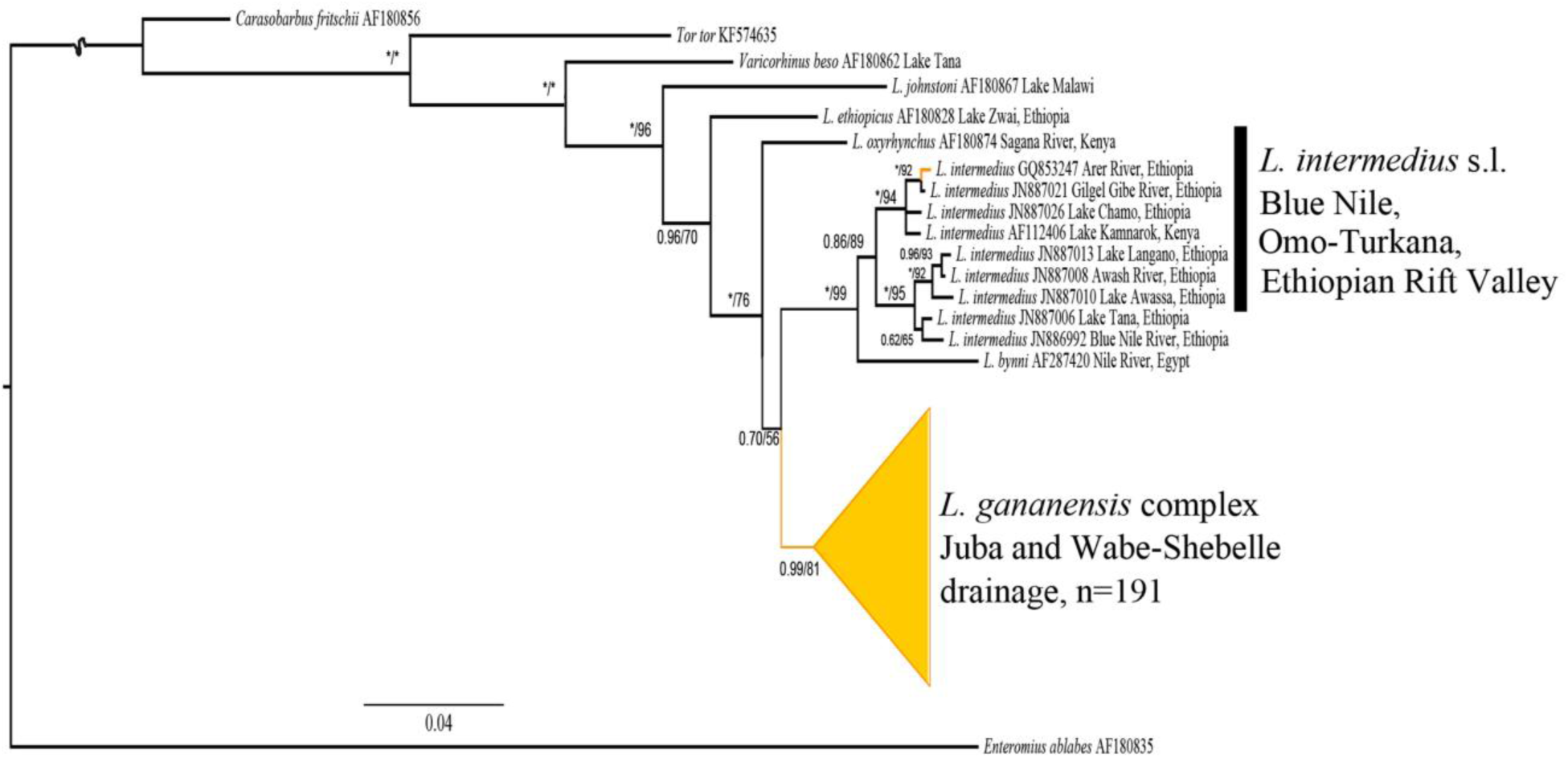
Bayesian Inference majority-rule consensus tree of relationships among the Ethiopian *Labeobarbus* barbs from all main drainages, including all unique haplotypes from the JWS drainages as well as *Labeobarbus* specimens from the surrounding drainages, based on cyt-*b* sequences. Bayesian posterior probabilities (on the left) and bootstrap values from ML analysis (on the right) above 0.5/50 are shown; asterisks represent posterior probabilities/bootstrap values of 1/100. Scale bar and branch lengths are given in expected substitutions per site. The nodes of JWS barbs were collapsed to a triangle, with the horizontal depth indicating the level of divergence within the node.

### 3.4. Relationships among barbs from the Genale and other localities

The BI tree constructed with the concatenated cyt-*b* + *d-loop* dataset (1664 bp) revealed a non-monophyletic origin of the Genale forms (Fig. 6). Polyphyly was decisively supported, particularly with the topological hypothesis testing of the ‘NEC’ form (log BF = 394.6). Since the tree topology does not fully resolve the relationships neither among only the Genale forms nor among Genale and other barb species, a haplotype network was also constructed with this dataset (Fig. 7).

**Fig. 6.**
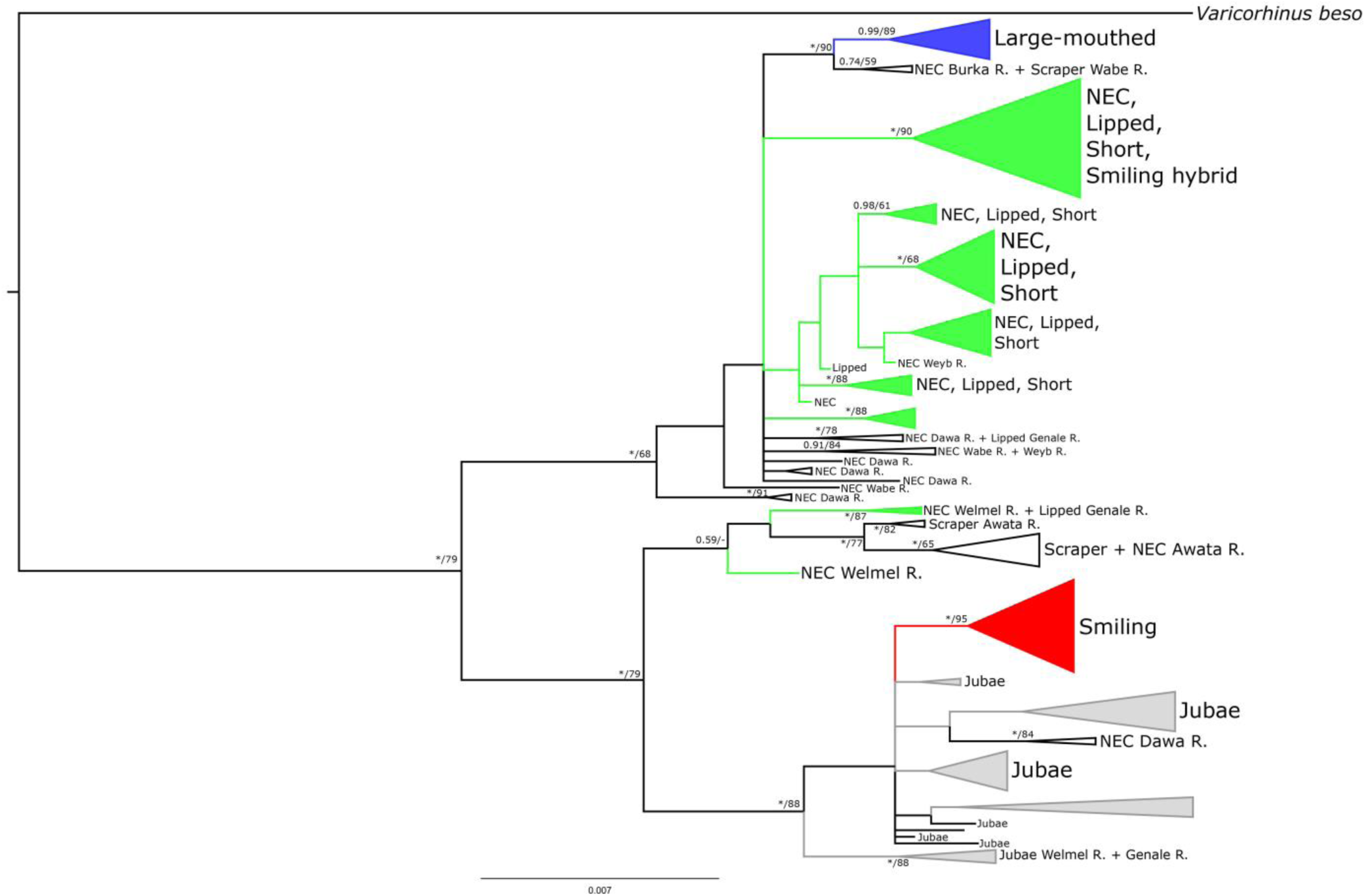
BI-tree of concatenated set (cyt-*b* + *dloop*) of the Genale forms and other JWS *Labeobarbus* barbs. Bayesian posterior probabilities (on the left) and bootstrap values from a ML analysis (on the right) above 0.5/50 are shown; asterisks represent posterior probabilities/bootstrap values of 1/100. Scale bar and branch lengths are in expected substitutions per site. Some nodes were collapsed to a triangle, with the horizontal depth indicating the level of divergence within the node. Genale barbs are colored by green (‘NEC’, ‘lipped’, and ‘short’), blue (‘large-mouthed’), red (‘smiling’), and grey (*L. jubae*).

**Fig. 7.**
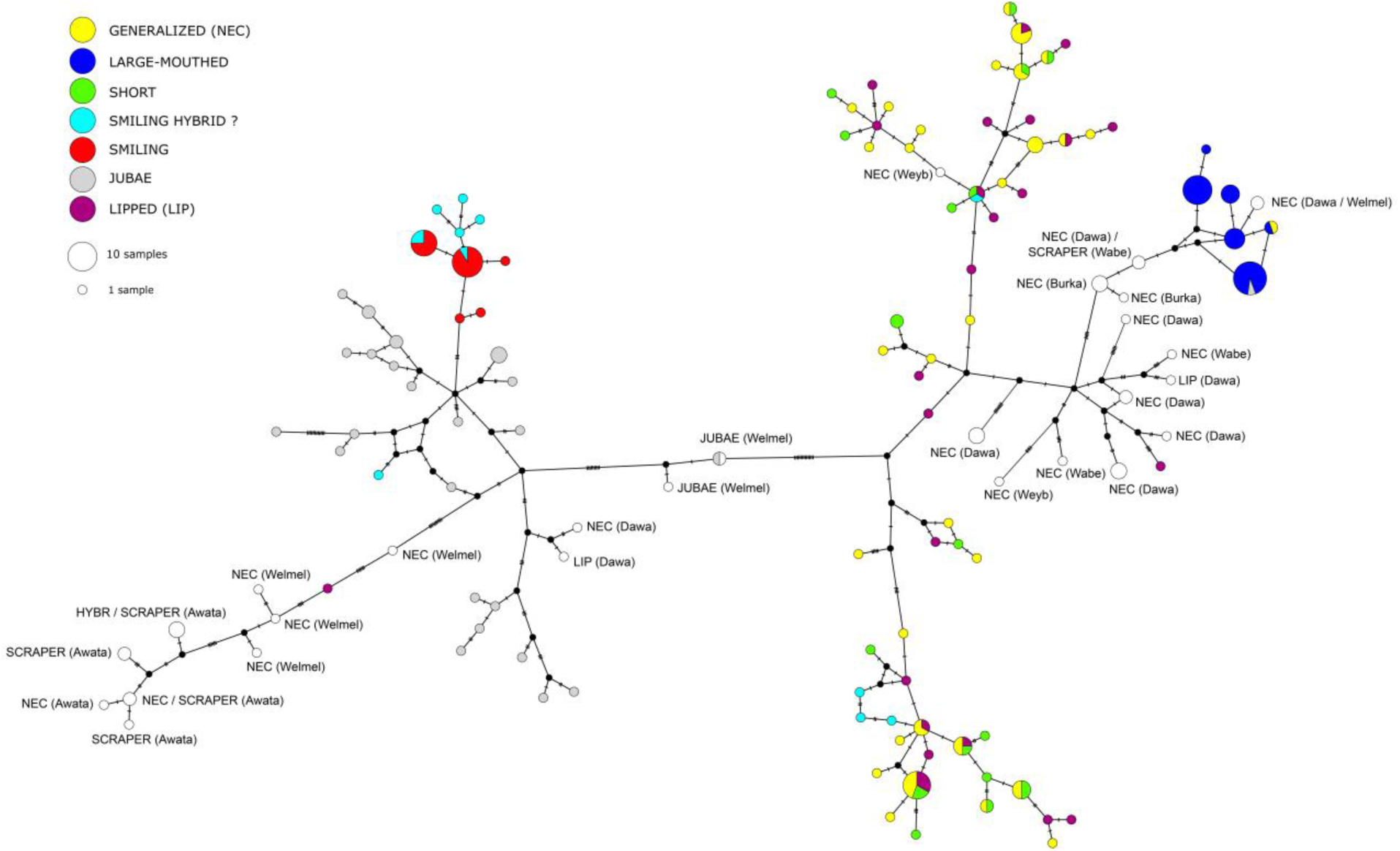
Median-joining haplotype network of the Genale assemblage and other barbs from JWS basin constructed on the basis of 221 *d-loop* sequences. Haplotypes of Genale barbs are colored, whereas those from other rivers are given as white circles with indication of their form (‘NEC’, ‘scraper’, ‘lipped’) and geographic allocation.

### 3.5. Haplotype network analysis of intra- and interspecific relationships of Genale barbs and barbs from other rivers in the JWS drainage

The haplotype network constructed with *d-loop* sequences revealed high diversity among Genale barbs and barbs from other rivers in the JWS drainage (Fig. 7). A total of 126 haplotypes were found (Supplementary Table S7). Intraspecific relationships among Genale forms suggest a common origin. ‘NEC’, ‘short’, and ‘lipped’ forms were represented in two clusters. Three specimens (two ‘lipped’ and one ‘NEC’) were assigned to different clusters (Fig. 7). One of the ‘lipped’ specimens occurs between ‘NEC’ barbs from the Welmel River (a tributary of the Genale River) and ‘NEC’ + Scraper barbs from the Awata River (a tributary of the Dawa River in the Juba drainage). The other ‘lipped’ specimen is placed with the ‘NEC’ form from the Dawa River (Fig. 7). It is important to note that although the Awata River is a tributary of the Dawa River, the ‘NEC’ forms from these localities are very distant in the network (min. 76 mutational steps; Fig. 7). In fact, they are genetically closer to populations from other rivers of the JWS than to each other. One Genale ‘NEC’ specimen shared the same haplotype with the Genale ‘large-mouthed’ form. Only one non-Genale ‘NEC’ haplotype was found within the Genale ‘NEC’— ‘short’— ‘lipped’ cluster. This was a specimen from the Weyb River, a tributary of the Juba River. Another Weyb ‘NEC’ specimen grouped together with Dawa ‘NEC’ specimens (Fig. 7). Individuals of the ‘short’ form shared haplotypes exclusively with Genale ‘NEC’ and ‘lipped’ forms (Fig. 7). The fact that this ‘short’ phenotype is only found in the Genale River suggests a local origin of this form. Only two ‘lipped’ individuals did not fall within the Genale clusters (Fig. 7). One was close to ‘NEC’ and ‘scraper’ specimens from the Awata River, and the other clustered with ‘NEC’, ‘lipped’ and ‘scraper’ forms from Rivers Dawa, Weyb, Burka, and Wabe.

The most genetically distinct group of Genale forms are the scrapers (particularly *L. jubae* and undescribed ‘smiling’ form; Fig. 7). Although they are only separated by three mutational steps, they do not share any haplotypes. However, one phenotypically intermediate specimen was found, which belonged to the *L. jubae* haplogroup.

The scraper *L. jubae* was described from the Genale River (Banister 1984), and is also found in other rivers of the Juba Basin (Levin et al. 2013). Most of these specimens were separated from the other Genale forms by at least 31 mutational steps (Fig. 7). In fact, Genale scraper forms were closer to ‘NEC’ and ‘lipped’ forms from other parts of the JWS drainage (Dawa basin) than to the Genale ‘NEC’—‘short’— ‘lipped’ cluster, providing evidence that the *L. jubae* scraper form originated outside of the Genale. Moreover, some specimens with distinct scraping features like *L. jubae* (the lower jaw is modified into a scraper) belonged to the ‘NEC’ and ‘lipped’ haplogroups in the Awata River (Juba basin) and Wabe River (Wabe-Shebelle basin).

The ‘large-mouthed’ form was represented by a compact haplogroup composed of six haplotypes (Fig. 7). This haplogroup is linked to Genale ‘NEC’—‘short’— ‘lipped’ clusters through other non-Genale ‘NEC’, ‘lipped’ and ‘scraper’ forms (Fig. 7). Two non-Genale ‘NEC’ specimens from the Burka River in the Wabe-Shebelle drainage clustered with the ‘large-mouthed’ haplogroup, with only one nucleotide substitution between them (Fig. 7). Another closely related pair of specimens (*i.e*. sharing the same haplotype) were ‘NEC’ and ‘scraper’ forms from the Wabe River (Fig. 7). One Genale ‘NEC’ specimen and one Genale *L. jubae* shared haplotypes with ‘large-mouthed’ specimens. This could be due to recent hybridization, since all other Genale ‘NEC’— ‘lipped’— ‘short’ and *L. jubae* individuals are clearly divergent from the ‘large-mouthed’ haplogroup (Fig. 7). Therefore, the ‘large-mouthed’ form likely originated from the Wabe-Shebelle drainage and later colonized a wider riverine network.

### 3.6. Genetic diversity, population structure and demography

With the exception of the ‘large-mouthed’ and ‘smiling’ forms, the Genale barbs are characterized by very high haplotype diversity (Supplementary Table S7). The lowest haplotype diversity was detected in the ‘large-mouthed’ and ‘smiling’ forms (0.76 and 0.74, respectively; Supplementary Table S7). The ‘smiling hybrid’ group had the highest nucleotide diversity (Supplementary Table S7). Tajima’s D did not differ significantly from zero in any of the Genale barbs, providing no evidence for selection (Table S7). Negative and significant Fu’s *Fs* values were observed in ‘NEC’, ‘lipped’ and *L. jubae* forms (Table S7), suggesting a recent population expansion or occurrence of genetic hitchhiking.

Low and non-significant *F*_ST_ values were detected between ‘NEC’ and ‘short’ (0.01), ‘NEC’ and ‘lipped’ (0.01), and ‘short’ and ‘lipped’ (0.01), suggesting a common gene pool among these forms (Table S8). *F*_*ST*_ values between other forms were very high, indicating limited gene flow. The highest *F*_ST_ values were detected between the ‘large-mouthed’ and ‘smiling’ forms (0.925), ‘NEC’ and ‘smiling’ (0.776), ‘short’ and ‘smiling’ (0.771), ‘large-Mouthed’ and *L. jubae* (0.762), and ‘lipped’ and ‘smiling’ (0.757) (Table S8).

## 4. Discussion

Within the Genale River, we have found an assemblage of six sympatric forms that appear to partition trophic resources using three different foraging strategies (*viz*. scraping, omnivory, piscivory). These specialized forms, including two diverse scraping forms, show morphological adaptations to the mode of feeding as evidenced by divergent stable isotope ratios. This assemblage appears to be a monophyletic group, divergent from other *Labeobarbus* populations from adjacent drainages. These results support the hypothesis that the Genale barbs assemblage is a species flock. However, the phylogeographic relationship between barbs from the Genale and other JWS drainages is complicated. We discuss these results in the context of adaptive radiations in fishes.

### 4.1. Trophic resource partitioning by sympatric Genale forms

Gill raker number and gut length are morphological features typically associated with trophic resource partitioning (Kramer and Bryant 1995; Horstkotte and Strecker 2005; Kahilainen et al. 2011). Hence, differences in gill raker counts likely reflect adaptation to different foraging strategies. With the addition of gut content and stable isotope analysis, we were able to provide further support for resource partitioning of the sympatric forms found in the Genale. For example, the gills of the ‘NEC’, ‘short’ and ‘lipped’ forms had cushions at the bottom of the pharyngeal rakers, which are saturated in taste buds, as also shown for a cyprinid *Cyprinus carpio* (Sibbing 1988). Similar cushions were described by Matthes (1963) in *Labeobarbus altianalis* (Boulenger, 1900), a species that is considered to be omnivorous. External taste buds direct searching the substrate for benthic food (detritus, insect larvae, *etc*.; Nagelkerke 1997). Hence, the three forms expressing these cushioned rakers saturated in taste buds are likely benthic omnivores – a finding also supported by gut length measurements and content analysis. Specifically, these three forms had long intestines – characteristic of omnivores – that were filled with a high proportion of detritus. Moreover, stable isotope analysis indicated that these forms occupy the lowest trophic level, as they had the lowest δ^15^N values, similar to sympatric detritophagous labeonins *Labeo* cf. *cylindricus* (mean δ^15^N = 11.2±0.1‰), bentophagous mormyrids *Mormyrus caschive* (mean δ^15^N = 11.6±0.2‰), and *L. intermedius* from Lake Awassa (10.8-11.8‰) that feed primarily on benthic molluscs (Desta et al. 2006). Notably, the slightly shorter gut and higher δ^15^N in the ‘lipped’ form may suggest a more benthophagous habitat than that occupied by the ‘NEC’ form, especially in light of the fact that enlarged lips are known to be an adaptation to foraging on benthos hidden between rock crevices on pebble and rock fragments (Matthes 1963). Nevertheless, there were very little differences between these three forms in all the phenotypic traits measured, except body shape, which was shorter in the ‘short’ form as a result of compressed vertebral bodies (Levin and Golubtsov 2015, see also Fig. S2). Whether this represents a genetic or plastic change, and whether it is adaptive or not remains to be explored.

Unlike the three forms described above, which had intermediate numbers of gill rakers, the two scraping forms (‘smiling’ and ‘*L. jubae*’) had the highest number of gill rakers that is consistent with knowledge on other specialized scrapers (e.g. Levin 2012). However, they differed in gut length, which was significantly longer in the ‘*L. jubae*’ form. This increased gut length is similar to that seen in the highly specialized algae scraper, *Varicorhinus beso*, which is the type species of the genus (6-7 SL in adults according to Levin 2012). In fact, the ‘*L. jubae*’ form was initially classified in the *Varicorhinus* genus, likely due to its morphological appearance of a specialized algae scraper (Banister 1984). Although the ‘smiling’ form demonstrates scraping features (wide mouth with reduced lips and a horny cutting edge on the lower jaw), its gut was shorter and also included benthic larvae of the family Simulidae attached to stones. Together with the higher values of δ^15^N, this indicates that the ‘smiling’ scraper has a mixed diet, as it likely grazes on algae as well as attached benthos. Hence, there were notable differences between the two scraping forms.

While having an increased number of gill rakers is beneficial for filtering algae and detritus, fewer gill rakers are needed when preying upon planktivorous fish. As such, divergence in gill raker numbers is often observed between omnivorous and piscivorous species (Hjelm et al. 2000; Alekseyev et al. 2002). Accordingly, the ‘large-mouthed’ form also displayed the fewest number of gill rakers, consistent with piscivory. Furthermore, this form had the shortest gut (roughly 2 SL). Although piscivory is relatively rare among cyprinids – due to the absence of oral teeth – some large piscivorous African barbs have been described in a species flock of *Labeobarbus* spp. from Lake Tana. In this case, the eight forms/species that were identified as piscivorous or predominantly piscivorous had shorter guts than the non-piscivorous forms (Nagelkerke 1997; Sibbing et al. 1998). Gut content and stable isotope analyses both confirmed the high trophic position of this form: the stomachs (when filled) contained juvenile fish, and the ^15^N was roughly 2.7‰ enriched relative to the ‘short’ form. In aquatic systems, the difference in δ^15^N values between trophic levels is usually about 2.3-3.0‰ (McCutchan et al. 2003; Vanderklift and Ponsard 2003), hence the ^15^N enrichment observed in the ‘large-mouthed’ form is indicative of a trophic shift. Moreover, the δ^15^N values of this form were similar to those of the predatory bagrid catfish *Bagrus urostigma* (mean δ^15^N = 14.1±0.2‰ – our unpublished data) inhabiting the same locality.

### 4.2. mtDNA divergence among the trophic forms of Genale barbs

Our phylogeographic analysis showed that groups defined by resource partitioning belong to different haplogroups. Specifically, the three omnivorous forms that showed high phenotypic similarities (*viz*. ‘NEC’, ‘lipped’, and ‘short’) also clustered together with little differentiation between pairs (*F*_ST_ < 0.01). Interestingly, these forms were represented in two relatively divergent clades/haplogroups, both composed of predominately Genale haplotypes. Most probably, such pattern reflects colonization of the Genale River by two divergent ancestral populations that experienced admixture in the Genale. Within each clade/haplogroup, there were several haplotypes shared among the three forms. This could suggest that they comprise a polymorphic population with phenotypic differences arising from plasticity. Alternatively, these three forms could be reproductively isolated, but if recent, this genetic divergence would not be evident in mtDNA.

Similarly, the ‘large-mouthed’ form formed an independent clade/haplogroup comprised primarily of large-mouthed individuals. This haplogroup was genetically similar to the ‘NEC’ forms from Dawa, Burka, Wabe and Weyb Rivers, suggesting a possible origin outside of the Genale River. Indeed, this ‘large-mouthed’ phenotype may have originated in the Wabe drainage and later migrated to the Genale River, however it is also possible that the Wabe ‘NEC’ phenotype first migrated to the Genale and later evolved into the ‘large-mouthed’ form. During our short visit to the Wabe River and its tributaries, we could only find a few ‘NEC’ specimens. In addition, we encountered one novel scraper form (arched, narrow, non-lipped mouth; lower jaw covered by a horny cutting edge). This previously undescribed scraper form in the Wabe basin could be a result of another independently evolved scraper mouth phenotype within the JWS drainage, but additional samples would be needed to test this possibility. In fact, the ‘large-mouthed’ phenotypes of *Labeobarbus* have been described as separate species, including *L. zaphiri* (Blue Nile, Ethiopia), *L. kiogae* (Victoria Nile and Lake Kyoga), *L. longirostris* (Lake Kyoga; Worthington 1929) or as undescribed forms of a polymorphic species (Golubtsov 1993; Dimmick et al. 2001; Mina et al. 2001; Golubtsov 2010; current study).

The two scraper forms were more genetically similar to populations composed of ‘NEC’ and scraper forms from Dawa basin tributaries than to ‘NEC’ and scraper forms from the Genale. mtDNA data suggests that the *L. jubae* scraper originated from a scraper population in the Dawa River, which is genetically similar to the sympatric ‘NEC’ population. Interestingly, the ‘smiling’ scraper form appears to have evolved from the Genale population of *L. jubae*, as it displays unique haplotypes.

### 4.3. Mechanisms of speciation in Genale barbs: allopatric vs. sympatric? Towards a concept of multi (yet mini) flock

When analyzed in a broad geographic context (with *cyt-b* only), all individuals of the *L. gananensis* complex from the JWS drainage formed a monophyletic group that was a sister clade to *Labeobarbus* spp. taxa from other distant drainages (e.g. Rift Valley, Nile, Omo-Turkana). This suggests that the adaptive radiation of the Genale labeobarbs originated within the JWS drainage, and this Genale assemblage might constitute a species flock. However, strict monophyly was not always supported when reference samples from other JWS localities were analyzed with the addition of *d-loop* sequences. Hence, it is likely that the assemblage of barbs in the Genale has both sympatric and allopatric origins. Specifically, the ‘NEC’, ‘lipped’ and ‘short’ forms likely diverged in sympatry within the Genale, whereas the scrapers and ‘large-mouthed’ forms originated from other regions in the JWS drainage (*i.e*. allopatric to the Genale, but sympatric with other forms elsewhere in the JWS drainage). Thus, the Genale labeobarbs assemblage is likely a result of several events of sympatric ecological divergence that occurred in geographically distant localities, with subsequent dissipation via riverine net. As the Genale River is centrally located in a broad riverine net of the JWS drainage, it could serve as a hub that facilitates migration from both eastern (Dawa) and western (Wabe-Shebelle) channels (Fig. 2).

Of course, we cannot exclude the possibility of secondary contact or secondary sympatry as additional promoters of phenotypic novelty. These events have been shown to help species overcome environmental challenges and adapt to novel environments during adaptive radiations (Mittelbach and Schemske 2015; Nichols et al. 2015; Meier et al. 2017). In addition, introgressive hybridization between phylogenetically distant Ethiopian *Labeobarbus* spp. is also possible, as recently reported (Levin and Golubtsov 2017).

A species flock is defined as a monophyletic assemblage of endemic forms or species geographically restricted to a particular area (Greenwood 1984; Schön and Martens 2004). Oliver (2016) expanded this concept to define a species metaflock: ‘a complex of two or more closely related species flocks (within the same subfamily or tribe), each at least presumptively monophyletic, that do not together make up a monophyletic group.’ However, the Genale barb assemblage does not directly adhere to this definition. We consider such an assemblage of sympatric species/forms composed of several independently originated mini-flocks (although being genetically closer to each other than to lineages outside the basin) as a *species multiflock*. This term might also apply to other cases, such as with other large African barbs widely distributed throughout the continent (Vreven et al. 2016, 2018).

### 4.4. The role of riverine environments in adaptive radiations

Until recently, tests for monophyly have generally not been conducted to satisfy the species flock concept in riverine systems (Berg 1914; Burnashev 1952; Aleksandrova and Kuznetsov 1967, 1968; Roberts 1998; Roberts and Khaironizam 2008) except some recent studies (e.g. Koblmüller et al. 2008; Schwarzer et al. 2011; Piálek et al. 2012, 2018). Compared to lacustrine environments, riverine conditions are usually thought to provide fewer opportunities for fish diversification. However, it has been suggested that some ancient riverine systems represent unique evolutionary hotspots with multiple intra-basin radiations (Roberts 1998; Glaubrecht and Köhler 2004; Feulner et al. 2007; Piálek et al. 2012; Bolotov et al. 2017). The JWS drainage basins have striking upstream canyons (see example in Fig. S3) that have been shaped by the succession of rifting events in the Miocene 18 to 11 mya (Wolfenden et al. 2004; Mège et al. 2015). Traces of antecedent rivers, including paleochannels currently filled with basalt lava of Miocene-Pleistocene age, suggest that the JWS basin is indeed ancient (Mège et al. 2015). Therefore, the dynamic geological history and the ancient origin of the JWS riverine net can explain the complex genetic structure of JWS labeobarbs. Impassable barriers for fish migration, such as 20 m high waterfalls (Fig. S4) still exist, hence gene flow between separate river sections is impeded. Moreover, it has been shown that waterfalls separating depauperate fish fauna from a richer fauna can induce trophic polymorphism in sympatry (Roberts and Khaironizam 2008; Golubtsov 2010). The species flock of *Labeobarbus* barbs in Lake Tana is a striking example that corroborates this suggestion. Lake Tana includes 15 endemic forms/species of *Labeobarbus* and eight other non-*Labeobarbus* species, which are physically isolated from the much richer fauna downstream of a very high waterfall (Tis-Isat; ca. 40 m high; Golubtsov 2010).

### 4.5. Adaptive radiation in the context of origin of Labeobarbus

Many reports on the mouth polymorphism of *Labeobarbus* spp. suggest that this is a common intrinsic feature of this group, rather than an exception. Such propensity to produce different trophic phenotypes is apparently explained by an allopolyploid origin of the evolutionary hexaploid lineage of *Labeobarbus* (Oellermann and Skelton 1990; Golubtsov and Krysanov 1993; Yang et al. 2015). The maternal lineage of the hexaploid *Labeobarbus* was from the tetraploid *Tor* lineage (Yang et al. 2015), widely distributed from Southeastern Asia via South Asia to the Middle East. Some representatives of the *Tor* lineage have a generalized mouth phenotype with moderately developed lips, while others have a phenotype with over-developed or hypertrophied ‘rubber lips’ (Borkenhagen 2014; Coad 2017). On the other hand, the paternal lineage was from the diploid *Cyprinion* (Yang et al. 2015), distributed in South Asia and the Middle East. Most species of this genus are specialized scrapers with a well-developed horny cutting edge on the lower jaw as well as other specialized features (Coad 2017). Therefore, a discrete trophic polymorphism of *Labeobarbus* is based on preexisting genetic templates inherited from its ancestors, and realized numerously as homoplasy in many isolated lineages distributed throughout Africa.

The genus *Labeobarbus* provides an excellent model system to study parallellism of adaptive radiations, and can help clarify the mechanisms by which mouth polymorphisms have aided avoidance of intraspecific competition, ultimately resulting in ecological speciation. Apparently, inherited genomic templates of the trophic polymorphism played a major role in the evolutionary success of this significantly diversified lineage (ca. 125 species sensu Vreven et al. 2016). Complex genomes that have experienced past hybridization may contribute to rapid adaptive radiation, as established in cichlids from African Great lakes (Meier et al. 2017; Irisarri et al. 2018). Ancient introgressions have also been detected in Arctic charr (*Salvelinus*; Lecaudey et al. 2018) which are classic examples of sympatric divergence in high altitudes (Hindar and Jonsson 1982; Alekseyev et al. 2002; Knudsen et al. 2006; Taylor 2016). Application of advanced molecular technologies to investigations of hexaploids can contribute deeper insights into the evolutionary history of *Labeobarbus* (Stobie et al. 2018).

## Supporting information

## Acknowledgments

This study was supported by Russian Science Foundation (grant 15-14-10020). The field operations during recent years were supported by Joint Ethiopian-Russian Biological Expedition (JERBE). We are very thankful to Fekadu Tefera and Genanaw Tesfaye as well as to JERBE coordinator, Andrey A. Darkov, for valuable support during field operations and logistics. We acknowledge Michael V. Mina and Sergey E. Cherenkov for their help in field operations. Sergey E. Cherenkov was kind to prepare photographs. We very appreciate to Michael V. Mina and Juha Merilä for valuable commentaries on first draft, and to Jacquelin DeFaveri for linguistical corrections.

## References

Akaike, H. 1973. Maximum likelihood identification of Gaussian autoregressive moving average models. Biometr. 60:255–265.

Aleksandrova, E. N., and V. V. Kuznetsov. 1967. Ecology of diadromus whitefishes in the Lena River during autumn-winter period. J. Ichthyol. 7:46–58.

Aleksandrova, E. N., and V. V. Kuznetsov. 1968. On intraspecies forms of Coregonus muksun (Pallas) from the Lena River. Bull. Mosc. Univ. Biol.-Soil. 1:28–37.

Alekseyev, S. S., V. P. Samusenok, A. N. Matveev, and M. Y. Pichugin. 2002. Diversification, sympatric speciation, and trophic polymorphism of Arctic charr, Salvelinus alpinus complex, in Transbaikalia. Env. Biol. Fish. 64:97–114.

Bandelt, H. J., P. Forster, and A. Röhl. 1999. Median-joining networks for inferring intraspecific phylogenies. Mol. Biol. Evol. 16:37–48.

Banister, K. E. 1973. A revision of the large Barbus (Pisces, Cyprinidae) of East and Central Africa: II. Studies of African Cyprinidae. Bull. Brit. Mus. Nat. Hist. Zool. 26:3–148.

Banister, K. E. 1984. Three new species of Varicorhinus (Pisces, Cyprinidae) from Africa. Bull. Brit. Mus. Nat. Hist. Zool. 47:273–282.

Barluenga, M., K. N. Stölting, W. Salzburger, M. Muschick, and A. Meyer. 2006. Sympatric speciation in Nicaraguan crater lake cichlid fish. Nature 439:719–723.

Berg, L. S. 1914. [Fishes (Marsipobranchii and Pisces). Vol 3. Ostariophysi. Fasc. 2.] Imper. Acad. Sci. Petrograd, pp. 335–704.

Beshera, K. A., and P. M. Harris. 2014. Mitochondrial DNA phylogeography of the Labeobarbus intermedius complex (Pisces, Cyprinidae) from Ethiopia. J. Fish Biol. 85:228–245.

Bolotov, I. N., I. V. Vikhrev, A. V. Kondakov, E. S. Konopleva, M. Y. Gofarov, O. V. Aksenova, and S. Tumpeesuwan. 2017. New taxa of freshwater mussels (Unionidae) from a species-rich but overlooked evolutionary hotspot in Southeast Asia. Sci. Rep. 7:11573.

Borkenhagen, K. 2014. A new genus and species of cyprinid fish (Actinopterygii, Cyprinidae) from the Arabian Peninsula, and its phylogenetic and zoogeographic affinities. Env. Biol Fish. 97:1179–1195.

Burnashev, M. S. 1952. Snow trouts of the Zeravshan River. Proceedings of Kishinev State University (Biol.). 4:111–125.

Castric, V., F. Bonney, and L. Bernatchez. 2001. Landscape structure and hierarchical genetic diversity in the brook charr, Salvelinus fontinalis. Evolution. 55:1016–1028.

Coad, B. 2017. Freshwater Fishes of Iran. Available at http://briancoad.com/Species%20Accounts/Contents%20new.htm. Accessed October 15, 2018.

de Graaf, M., Eshete Dejen, J. W. M. Osse, and F. A. Sibbing. 2008. Adaptive radiation of Lake Tana’s (Ethiopia) Labeobarbus species flock (Pisces, Cyprinidae). Mar. Freshwater Res. 59:391–407.

de Graaf, M., H. J. Megens, J. Samallo, and F. Sibbing. 2010. Preliminary insight into the age and origin of the Labeobarbus fish species flock from Lake Tana (Ethiopia) using the mtDNA cytochrome b gene. Mol. Phylogen. Evol. 54:336–343.

Desta, Z., R. Borgstrøm, B. O. Rosseland, and Z. Gebre‐Mariam. 2006. Major difference in mercury concentrations of the African big barb, Barbus intermedius (R.) due to shifts in trophic position. Ecol. Freshwat. Fish. 15:532–543.

Dgebuadze, Y. Y. 1995. The land/inland-water ecotones and fish population of Lake Valley (West Mongolia). Hydrobiol. 303:235–245.

Dimmick, W. W., P. B. Berendzen, and A. S. Golubtsov. 2001. Genetic comparison of three Barbus (Cyprinidae) morphotypes from the Genale River, Ethiopia. Copeia. 2001:1123–1129.

Excoffier, L., and H. Lisher. 2010. Arlequin suite ver. 3.5: a new series of programs to perform population genetics analyses under Linux and Windows. Mol. Ecol. Res. 10:564–567.

Elmer, K. R., T. K. Lehtonen, A. F. Kautt, C. Harrod, and A. Meyer. 2010. Rapid sympatric ecological differentiation of crater lake cichlid fishes within historic times. BMC Biol. 8:60.

Feulner, P. G. D., F. Kirschbaum, V. Mamonekene, V. Ketmaier, and R. Tiedemann. 2007. Adaptive radiation in African weakly electric fish (Teleostei: Mormyridae: Campylomormyrus): a combined molecular and morphological approach. J. Evol. Biol. 20:403–414.

Franchini, P., C. Fruciano, M. L. Spreitzer, J. C. Jones, K. R. Elmer, F. Henning, and A. Meyer. 2014. Genomic architecture of ecologically divergent body shape in a pair of sympatric crater lake cichlid fishes. Mol. Ecol. 23:1828–1845.

Fu, Y. X. 1997. Statistical tests of neutrality of mutations against population growth, hitchhiking and background selection. Genetics. 147:915–925.

Glaubrecht, M., and F. Köhler. 2004. Radiating in a river: systematics, molecular genetics and morphological differentiation of viviparous freshwater gastropods endemic to the Kaek River, central Thailand (Cerithioidea, Pachychilidae). Biol. J. Linn. Soc. 82:275–311.

Glez-Peña, D., D. Gómez-Blanco, M. Reboiro-Jato, F. Fdez-Riverola, and D. Posada. 2010. ALTER: program-oriented format conversion of DNA and protein alignments. Nucl. Acids Res. Web Server issue. ISSN: 0305–1048 http://dx.doi.org/10.1093/nar/gkq321

Golubtsov, A. S., 1993. Biogeographie des “grands Barbus” d’Éthiopie avec référence spéciale à des formes a status taxinomiques incertains. Cah. Ethol. 13:227–230.

Golubtsov, A. S. 2010. Species flocks of fishes in rivers and lakes: sympatric divergence in faunistically depauperate fish communities as especial modus of evolution. In: D. S. Pavlov, Y. Y. Dgebuadze, and M. I. Shatunovskiy (eds.) Current problems of ichthyology. To 100-jubelee of G.V. Nikolsky, KMK, Moscow, 96–123 [in Russian].

Golubtsov, A. S., and E. Y. Krysanov. 1993. Karyological study of some cyprinid species from Ethiopia. The ploidy differences between large and small Barbus of Africa. J. Fish Biol. 42: 445–455.

Gordeeva, N. V., S. S. Alekseyev, A. N. Matveev, and V. P. Samusenok. 2014. Parallel evolutionary divergence in Arctic char Salvelinus alpinus complex from Transbaikalia: variation in differentiation degree and segregation of genetic diversity among sympatric forms. Can. J. Fisher. Aquat. Sci. 72:96–115.

Greenwood, P. H. 1974. The cichlid fishes of Lake Victoria, East Africa: the biology and evolution of a species flock. Bull. Brit. Mus. Nat. Hist. Zool. Suppl. 6:1–134.

Greenwood, P. H. 1984. African cichlids and evolutionary theories. In I. Kornfield, and A. A. Echelle (eds.). Evolution of Fish Species Flocks. University of Maine Press, Orono, pp. 141–154.

Evolution of fish species flocks.

Herre, A. W. 1933. The fishes of Lake Lanao: a problem in evolution. Am. Soc. Nat. 67:154–162.

Hindar, K., and B. Jonsson. 1982. Habitat and food segregation of dwarf and normal Arctic charr (Salvelinus alpinus) from Vangsvatnet Lake, western Norway. Can. J. Fisher. Aquat. Sci. 39:1030–1045.

Horstkotte, J., and U. Strecker. 2005. Trophic differentiation in the phylogenetically young Cyprinodon species flock (Cyprinodontidae, Teleostei) from Laguna Chichancanab (Mexico). Biol. J. Linn. Soc. 85:125–134.

Irisarri, I., P. Singh, S. Koblmüller, J. Torres-Dowdall, F. Henning, P. Franchini, C. Fischer, A. R. Lemmon, E. M. Lemmon, G. G. Thallinger, C. Sturmbauer, and A. Meyer. 2018. Phylogenomics uncovers early hybridization and adaptive loci shaping the radiation of Lake Tanganyika cichlid fishes. Nat. Com. 9:3159.

Kahilainen, K. K., A. Siwertsson, K. Ø. Gjelland, R. Knudsen, T. Bøhn, and P. A. Amundsen. 2011. The role of gill raker number variability in adaptive radiation of coregonid fish. Evol. Ecol. 25:573–588.

Knudsen, R., A. Klemetsen, P. A. Amundsen, and B. Hermansen. 2006. Incipient speciation through niche expansion: an example from the Arctic charr in a subarctic lake. Proc. Royal Soc. Lond. B Biol. Sci. 273:2291–2298.

Kramer, D. L., and M. J. Bryant. 1995. Intestine length in the fishes of a tropical stream: 2. Relationships to diet—the long and short of a convoluted issue. Env. Biol. Fish. 42:129–141.

Koblmüller, S., K. M. Sefc, N. Duftner, C. Katongo, T. Tomljanovic, and C. Sturmbauer. 2008. A single mitochondrial haplotype and nuclear genetic differentiation in sympatric colour morphs of a riverine cichlid fish. J. Evol. Biol. 21:362–367.

Kondrashov, A. S., and M. V. Mina. 1986. Sympatric speciation: when is it possible? Biol. J. Linn. Soc. 27:201–223.

Kontula, T., S. V. Kirilchik, and R. Väinölä. 2003. Endemic diversification of the monophyletic cottoid fish species flock in Lake Baikal explored with mtDNA sequencing. Mol. Phylogen. Evol. 27:143–155.

Kornfield, I., and A. A. Echelle (eds.). 1984. Evolution of Fish Species Flocks. University of Maine Press, Orono.

Lanfear, R., B. Calcott, S. Y. W. Ho, and S. Guindon. 2012. PartitionFinder: combined selection of partitioning schemes and substitution models for phylogenetic analyses. Mol. Biol. Evol. 29:1695–1701.

Laporte, M., S. M. Rogers, A. M. Dion-Côté, E. Normandeau, P. A. Gagnaire, A. C. Dalziel, J. Chebib, and L. Bernatchez. 2015. RAD-QTL Mapping Reveals Both Genome-Level Parallelism and Different Genetic Architecture Underlying the Evolution of Body Shape in Lake Whitefish (Coregonus clupeaformis) Species Pairs. G3: Genes| Genomes| Genetics. g3:115.

Lecaudey, L. A., U. K. Schliewen, A. G. Osinov, E. B. Taylor, L. Bernatchez, and S. J. Weiss. 2018. Inferring phylogenetic structure, hybridization and divergence times within Salmoninae (Teleostei: Salmonidae) using RAD-sequencing. Mol. Phylogen. Evol. 124:82–99.

Leigh, J. W., and D. Bryant. 2015. Popart: full‐feature software for haplotype network construction. Methods Ecol. Evol. 6:1110–1116.

Levin, B. A. 2012. New data on morphology of the African scraping feeder Varicorhinus beso (Osteichthyes: Cyprinidae) with the special reference to specialized traits. J Ichthyol. 52:908–923.

Levin, B. A., and A. S. Golubtsov. 2015. Massive emergence of “short tail” deformity in the Labeobarbus species flock from the Genale River, Ethiopia. Abstr. Fourth Meeting “Interdisciplinary Approaches in Fish Skeletal Biology”, Portugal, Tavira.

Levin, B. A., and A. S. Golubtsov. 2017. An evidence of past introgressive hybridization between Labeobarbus ethiopicus and L. intermedius in the Ethiopian Rift Valley, East Africa. Ethiop. Biol. J. 16(Supplementary):45–60.

Levin, B. A., A. S. Golubtsov, Y. Y. Dgebuadze, and N. S. Mugue. 2013. New evidence of homoplasy within the African genus Varicorhinus (Cyprinidae): an independent origin of specialized scraping forms in the adjacent drainage systems of Ethiopia inferred from mtDNA analysis. Afr. Zool. 48:400–406.

Librado, P. and J. Rozas. 2009. DnaSP v5: a software for comprehensive analysis of DNA polymorphism data. Bioinform. 25:1451–1452.

Machado‐Schiaffino, G., F. Henning, and A. Meyer. 2014. Species‐specific differences in adaptive phenotypic plasticity in an ecologically relevant trophic trait: hypertrophic lips in Midas cichlid fishes. Evolution. 68:2086–2091.

Matthes, H. 1963. A comparative study of the feeding mechanisms of some African Cyprinidae (Pisces, Cypriniformes). Bijdragen tot de Dierkunde. 3:3–35.

McCutchan, J. H., W. M. Lewis, C. Kendall, and C. C. McGrath. 2003. Variation in trophic shift for stable isotope ratios of carbon, nitrogen, and sulfur. Oikos. 102:378–390.

McKinnon, J. S., and H. D. Rundle. 2002. Speciation in nature: the threespine stickleback model systems. Trends Ecol. Evol. 17:480–488.

Mège, D., P. Purcell, S. Pochat, and T. Guidat. 2015. The landscape and landforms of the Ogaden, Southeast Ethiopia. In Landscapes and Landforms of Ethiopia (pp. 323–348). Springer, Dordrecht.

Meier, J. I., D. A. Marques, S. Mwaiko, C. E. Wagner, L. Excoffier, and O. Seehausen. 2017. Ancient hybridization fuels rapid cichlid fish adaptive radiations. Nat. Com. 8:14363.

Meyer, A., T. D. Kocher, P. Basasibwaki, and A. C. Wilson. 1990. Monophyletic origin of Lake Victoria cichlid fishes suggested by mitochondrial DNA sequences. Nature. 347:550–553.

Meyer, A., J. M. Morrissey, and M. Schartl. 1994. Recurrent origin of a sexually selected trait in Xiphophorus fishes inferred from a molecular phylogeny. Nature. 368:539–542.

Mina, M. V., A. N. Mironovsky, and Y. Y. Dgebuadze. 1996. Lake Tana large barbs: phenetics, growth and diversification. J. Fish Biol. 48:383–404.

Mina, M.V., A. N. Mironovsky, A. S. Golubtsov, and Y. Y. Dgebuadze. 1998. ‘Barbus’ intermedius Species Flock in Lake Tana (Ethiopia). II. Morphological Diversity of ‘Large Barbs’ from Lake Tana and Neighbouring Areas: Homoplasies or Synapomorphies? Ital. J. Zool. 65:9–14.

Mina, M. V., A. N. Mironovsky, and D. Golani. 2001. Consequences and modes of morphological diversification of East African and Eurasian barbins (genera Barbus, Varicorhinus and Capoeta) with particular reference to Barbus intermedius complex. Env. Biol. Fish. 61:241–252.

Mittelbach, G. G., and D. W. Schemske. 2015. Ecological and evolutionary perspectives on community assembly. Trends Ecol. Evol. 30:241–247.

Myers, G. S. 1960. The endemic fish fauna of Lake Lanao, and the evolution of higher taxonomic categories. Evolution. 14:323–333.

Nagelkerke, L. 1997. The barbs of Lake Tana, Ethiopia: morphological diversity and its implications for taxonomy, trophic resource partitioning, and fisheries. PhD dissertation thesis. Wageningen Univ

Nagelkerke, L. A., F. A. Sibbing, J. G. van den Boogaart, E. H. Lammens, and J. W. Osse. 1994. The barbs (Barbus spp.) of Lake Tana: a forgotten species flock? Env. Biol. Fish. 39:1–22.

Nagelkerke, L. A. J., K. M. Leon‐Kloosterziel, H. J. Megens, M. De Graaf, O. E. Diekmann, and F. A. Sibbing. 2015. Shallow genetic divergence and species delineations in the endemic Labeobarbus species flock of Lake Tana, Ethiopia. J. Fish Biol. 87:1191–1208.

Nichols, P., M. J. Genner, C. Van Oosterhout, A. Smith, P. Parsons, H. Sungani, J. Swanstrom, and D. A. Joyce. 2015. Secondary contact seeds phenotypic novelty in cichlid fishes. Proc. Royal Soc. Lond. B Biol. Sci. 282:20142272.

Oellermann, L. K., and P. H. Skelton. 1990. Hexaploidy in yellowfish species (Barbus, Pisces, Cyprinidae) from southern Africa. J. Fish Biol. 37:105–115.

Oliver, M. K. 2016. A Kink in the Line: Does a Unique Lateral-Line Peculiarity Really Characterize Lake Malaŵi's Huge Haplochromine Species Flock (Teleostei: Cichlidae)? Bull. Peab. Mus. Nat. Hist. 57:3–20.

Østbye, K., P. A. Amundsen, L. Bernatchez, A. Klemetsen, R. Knudsen, R. Kristoffersen, T. F. Næsje, and K. Hindar. 2006. Parallel evolution of ecomorphological traits in the European whitefish Coregonus lavaretus (L.) species complex during postglacial times. Mol. Ecol. 15:3983–4001.

Palumbi, S. R. 1996. Nucleic acids II: the polymerase chain reaction. In: D. M. Hillis, C. Moritz, B. K. Mable (Eds.). Molecular Systematics. Sinauer Associates, Sunderland, MA, pp. 205–247.

Perdices, A., and I. Doadrio. 2001. The molecular systematics and biogeography of the European cobitids based on mitochondrial DNA sequences. Mol. Phylogen. Evol. 19:468–478.

Piálek, L., O. Říčan, J. Casciotta, A. Almirón, and J. Zrzavý. 2012. Multilocus phylogeny of Crenicichla (Teleostei: Cichlidae), with biogeography of the C. lacustris group: species flocks as a model for sympatric speciation in rivers. Mol. Phylogen. Evol. 62:46–61.

Piálek, L., E. Burress, K. Dragová, A. Almirón, J. Casciotta, and O. Říčan. 2018 (in press). Phylogenomics of pike cichlids (Cichlidae: Crenicichla) of the C. mandelburgeri species complex: rapid ecological speciation in the Iguazú River and high endemism in the Middle Paraná basin. Hydrobiol. 1–21.

Post, D. M., C. A. Layman, D. A. Arrington, G. Takimoto, J. Quattrochi, and C. G. Montana. 2007. Getting to the fat of the matter: models, methods and assumptions for dealing with lipids in stable isotope analyses. Oecologia. 152:179–189.

Rambaut, A., M. A. Suchard, D. Xie, A. J. Drummond. 2014. Tracer v1.6. http://beast.bio.ed.ac.uk/Tracer (accessed 3 August 2017).

Ramos-Onsins, S. E., and J. Rozas. 2002. Statistical properties of new neutrality tests against population growth. Mol. Biol. Evol. 19:2092–2100.

Roberts, T. R. 1998. Review of the tropical Asian cyprinid fish genus Poropuntius, with descriptions of new species and trophic morphs. Nat. Hist. Bull. Siam Soc. 46:105–135.

Roberts, T. R., and M. Z. Khaironizam. 2009. Trophic polymorphism in the Malaysian fish Neolissochilus soroides and other old world barbs (Teleostei, Cyprinidae). Nat. Hist. Bull. Siam Soc. 56:25–53.

Ronquist, F., M. Teslenko, P. van der Mark, D. L. Ayres, A. Darling, S. Höhna, B. Larget, L. Liu, M. A. Suchard, and J. P. Huelsenbeck. 2012. MrBayes 3.2: efficient Bayesian phylogenetic inference and model choice across a large model space. Syst. Biol. 61:539–542.

Salzburger, W., A. Meyer, S. Baric, E. Verheyen, and C. Sturmbauer. 2002. Phylogeny of the Lake Tanganyika cichlid species flock and its relationship to the Central and East African haplochromine cichlid fish faunas. Syst. Biol. 51:113–135.

Schliewen, U. K., D. Tautz, and S. Pääbo. 1994. Sympatric speciation suggested by monophyly of crater lake cichlids. Nature. 368:629–632.

Schluter, D. 1996. Ecological causes of adaptive radiation. Amer. Nat. 148:40–64.

Schluter, D. 2000. The ecology of adaptive radiation. Oxford University Press, NY.

Schön, I., and K. Martens. 2004. Adaptive, pre-adaptive and non-adaptive components of radiations in ancient lakes: a review. Organ. Divers. Evol. 4:137–156.

Schwarzer, J., B. Misof, S. N. Ifuta, and U. K. Schliewen. 2011. Time and origin of cichlid colonization of the lower Congo rapids. PLoS One. 6:e22380.

Seehausen, O., E. Koetsier, M. V. Schneider, L. J. Chapman, C. A. Chapman, M. E. Knight, G. F. Turner, J. J. M. van Alphen, and R. Bills. 2003. Nuclear markers reveal unexpected genetic variation and a Congolese-Nilotic origin of the Lake Victoria cichlid species flock. Proc. Royal Soc. Lond. B Biol. Sci. 270:129–137.

Seehausen, O., and C. E. Wagner. 2014. Speciation in freshwater fishes. Ann. Rev. Ecol. Evol. Syst. 45:621–651.

Sibbing, F. A. 1988. Specializations and limitations in the utilization of food resources by the carp, Cyprinus carpio: a study of oral food processing. Env. Biol. Fish. 22:161–178.

Sibbing, F. A., L. A. Nagelkerke, R. J. Stet, and J. W. Osse. 1998. Speciation of endemic Lake Tana barbs (Cyprinidae, Ethiopia) driven by trophic resource partitioning: a molecular and ecomorphological approach. Aquat. Ecol. 32:217–227.

Stamatakis, A. 2007. RAxML-VI-HPC: maximum likelihood-based phylogenetic analyses with thousands of taxa and mixed models. Bioinform. 22:2688–2690.

Stamatakis, A., P. Hoover, and J. Rougemont. 2008. A rapid bootstrap algorithm for the RAxML web servers. Syst. Biol. 575:758–771.

Stobie, C. S., C. J. Oosthuizen, M. J. Cunningham, and P. Bloomer. 2018. Exploring the phylogeography of a hexaploid freshwater fish by RAD sequencing. Ecol. Evol. 8:2326–2342.

Sullivan, J. P., S. Lavoué, and C. D. Hopkins. 2002. Discovery and phylogenetic analysis of a riverine species flock of African electric fishes (Mormyridae: Teleostei). Evolution. 56:597–616.

Tajima, F. 1989. Statistical method for testing the neutral mutation hypothesis by DNA polymorphism. Genetics. 123:585–595.

Tamura, K., G. Stecher, D. Peterson, A. Filipski, and S. Kumar. 2013). MEGA6: molecular evolutionary genetics analysis version 6.0. Mol. Biol. Evol. 30:2725–2729.

Taylor, E. B. 2016. The Arctic char (Salvelinus alpinus) “complex” in North America revisited. Hydrobiol. 783:283–293.

Thompson, J. D., D. G. Higgins, and T. J. Gibson. 1994. CLUSTAL W: improving the sensitivity of progressive multiple sequence alignment through sequence weighting, position-specific gap penalties and weight matrix choice. Nucl. Acids Res. 22:4673–4680.

Vanderklift, M. A., and S. Ponsard. 2003. Sources of variation in consumer-diet δ 15 N enrichment: a meta-analysis. Oecologia. 136:169–182.

Vreven, E. J., T. Musschoot, E. Decru, S. Wamuini Lunkayilakio, K. Obiero, A. F. Cerwenka, and U. K. Schliewen. 2018. The complex origins of mouth polymorphism in the Labeobarbus (Cypriniformes: Cyprinidae) of the Inkisi River basin (Lower Congo, DRC, Africa): insights from an integrative approach. Zool. J. Linn. Soc. zly049.

Vreven, E. J., T. Musschoot, J. Snoeks, and U. K. Schliewen. 2016. The African hexaploid Torini (Cypriniformes: Cyprinidae): review of a tumultuous history. Zool. J. Linn. Soc. 177:231–305.

Weir, B. S., and C. C. Cockerham. 1984. Estimating F‐statistics for the analysis of population structure. Evolution. 38:1358–1370.

Wolfenden, E., C. Ebinger, G. Yirgu, A. Deino, and D. Ayalew. 2004. Evolution of the northern Main Ethiopian rift: birth of a triple junction. Earth Planet. Sci. Let. 224:213–228.

Worthington, E. B. 1929. New Species of Fish from the Albert Nyanza and Lake Kioga. J. Zool. 99:429–440.

Yang, L., T. Sado, M. V. Hirt, E. Pasco-Viel, M. Arunachalam, J. Li, X. Wang, J. Freyhof, K. Saitoh, A. M. Simons, M. Miya, S. He, and R. L. Mayden. 2015. Phylogeny and polyploidy: resolving the classification of cyprinine fishes (Teleostei: Cypriniformes). Mol. Phylogen. Evol. 85:97–116.

