## Supplementary material for "Adaptive radiation of barbs of the genus *Labeobarbus* (Cyprinidae) in the East African river"

Supplementary Tables

Table S1. Species names, sampling localities, and GenBank accession numbers of material studied.

| **Species/forms** | **Sample size for genetics / stable isotope analysis** | **GenBank Acc. Nos., cyt-*b*** | **GenBank Acc. Nos., *d-loop*** | **Source of DNA sequences*** |
|---|---|---|---|---|
| **(1) Genale River, Juba R. basin, Ethiopia, 05°42'08.54"N, 39°32'39.30"E, altitude 1124 m** | | | | |
| *Labeobarbus gananensis*, 'NEC' form | 34/21 | MK001131-165 | MK001305-35 | This study |
| | | - | AF352842-51 JQ701772-73 | 1; 2 |
| *L. gananensis,* 'lipped' form | 18/14 | MK001166-181 | MK001288-304 | This study |
| | | - | AF352852-60 | 1 |
| *L. gananensis,* 'short' form | 20/20 | MK001056-74 | MK001336-54 | This study |
| | | - | JQ701771 | 2 |
| *L. jubae* | 28/14 | MK001092-118 | MK001268-87 | This study |
| | | - | JQ701763-70 | 2 |
| *Labeobarbus* sp.1 'smiling' form | 19/14 | MK001075-91 | MK001237-55 | This study |
| *Labeobarbus* sp.2 'large-mouthed' form | 29/26 | MK001027-55 | AF352861-64; JQ701756-57; MK001210-36 | 1; 2; this study |
| Putative hybrid between *L. gananensis* 'NEC' and 'smiling' form | 12/7 | MK001119-130 | MK001256-67 | This study |
| **(2) Welmel R., tributary of Genale R., Juba basin, Ethiopia, 06°13'34.00"N, 39°49'10.40"E, altitude 1014 m** | | | | |

| | | | | |
|---|---|---|---|---|
| *L. gananensis*, 'NEC' form | 5 | MK001207-09 | MK001388-92 | This study |
| *L. jubae* | 2 | MK001025-26 | JQ701761-62 | 2; this study |
| **(3) Dawa R., Juba basin, Ethiopia, 05°19'42"N, 38°47'13"E, altitude 1165 m** | | | | |
| *L. gananensis*, 'NEC' form | 13 | MK001186-95 | MK001365-77 | This study |
| *L. gananensis*, 'lipped' form | 2 | MK001185 | MK001364, MK001378 | |
| **(4) Awata R., tributary of Dawa R., Juba basin, Ethiopia, 05°47'07"N, 38°55'46"E, altitude 1632 m** | | | | |
| *L. gananensis*, 'NEC' form | 2 | MK001200-01 | MK001357-58 | This study |
| *L. jubae* | 9 | MK001196-99; MK001202-06 | JQ701758-60; MK001355-56; MK001359-63 | 2; this study |
| **(5) Weyb R., Juba basin, Ethiopia, 06°53'54"N, 40°51'06"E, altitude 1203 m** | | | | |
| *L. gananensis*, 'NEC' form | 2 | MK001182-83 | MK001382-83 | This study |
| **(6) Burka R., tributary of Wabe-Shebelle R., Ethiopia, 07°29'47"N, 40°58'53"E, altitude 1266 m** | | | | |
| *L. gananensis*, 'NEC' form | 4 | MK001184 | MK001384-87 | This study |
| **(7) Wabe-Shebelle R., Ethiopia, 07°27'17"N, 40°51'06"E, altitude 1076 m** | | | | |
| *L. gananensis*, 'NEC' form | 2 | MK015651-52 | MK001380-81 | This study |
| *L. gananensis*, 'scraping' form** | 1 | MK015650 | MK001379 | This study |

* 1 – Dimmick et al., 2001; 2 – Levin et al., 2013. ** Body shape and other features of this individual were similar to the 'NEC' form except for the presence of a weakly developed horny cutting edge on the lower jaw.

Table S2. Comparative cyt-*b* sequences mined from GenBank

| **Species** | **Locality** | **GenBank Acc. Nos., cyt-*b*** | **Source of DNA sequences** |
|---|---|---|---|
| *Labeobarbus bynni* | Nile R., Egypt | AF287420 | Machordom & Doadrio, 2001. |
| *L. ethiopicus* | Zwai Lake basin, Ethiopia | AF180828 | Tsigenopoulos et al., 2002 |
| *L. gananensis* | Genale River, Ethiopia | JN887036-38 | Beshera & Harris, 2014 |
| *L. intermedius* | Lake Kamnarok, Kenya | AF112406 | Tsigenopoulos & Berrebi, 2000 |
| *L. intermedius* | Blue Nile River, Ethiopia | JN886992 | Beshera & Harris, 2014 |
| *L. intermedius* | Lake Tana, Ethiopia | JN887006 | Beshera & Harris, 2014 |
| *L. intermedius* | Awash River, Ethiopia | JN887008 | Beshera & Harris, 2014 |
| *L. intermedius* | Lake Awassa, Ethiopia | JN887010 | Beshera & Harris, 2014 |
| *L. intermedius* | Lake Langano, Ethiopia | JN887013 | Beshera & Harris, 2014 |
| *L. intermedius* | Gilgel Gibe River, Omo-Turkana system, Ethiopia | JN887021 | Beshera & Harris, 2014 |
| *L. intermedius* | Lake Chamo, Ethiopia | JN887026 | Beshera & Harris, 2014 |
| *L. intermedius* | Arer River, Wabe-Shebelle system | GQ853247 | de Graaf et al., 2010 |
| *L. johnstonii* | Lake Malawi, Malawi | AF180867 | Tsigenopoulos et al., 1999 (unpubl.) |
| *L. oxyrhynchus* | Sagana River, Kenya | AF180874 | Tsigenopoulos et al., 1999 (unpubl.) |

| | | | |
|---|---|---|---|
| *Varicorhinus beso* | Lake Tana, Ethiopia | AF180862 | Durand et al., 2002 |

Literature cited:


Beshera, K. A., & Harris, P. M. (2014). Mitochondrial DNA phylogeography of the Labeobarbus intermedius complex (Pisces, Cyprinidae) from Ethiopia. *Journal of fish biology*, *85*(2), 228-245.

de Graaf, M., Megens, H. J., Samallo, J., & Sibbing, F. (2010). Preliminary insight into the age and origin of the Labeobarbus fish species flock from Lake Tana (Ethiopia) using the mtDNA cytochrome b gene. *Molecular Phylogenetics and Evolution*, *54*(2), 336-343.

Durand, J. D., Tsigenopoulos, C. S., Ünlü, E., & Berrebi, P. (2002). Phylogeny and biogeography of the family Cyprinidae in the Middle East inferred from cytochrome b DNA—evolutionary significance of this region. *Molecular Phylogenetics and Evolution*, *22*(1), 91-100.

Machordom, A., & Doadrio, I. (2001). Evolutionary history and speciation modes in the cyprinid genus Barbus. *Proceedings of the Royal Society of London B: Biological Sciences*, *268*(1473), 1297-1306.

Tsigenopoulos, C. S., & Berrebi, P. (2000). Molecular phylogeny of North Mediterranean freshwater barbs (genus Barbus: Cyprinidae) inferred from cytochrome b sequences: biogeographic and systematic implications. *Molecular Phylogenetics and Evolution*, *14*(2), 165-179.

Tsigenopoulos, C. S., Ráb, P., Naran, D., & Berrebi, P. (2002). Multiple origins of polyploidy in the phylogeny of southern African barbs (Cyprinidae) as inferred from mtDNA markers. *Heredity*, *88*(6), 466.

Table S3.

Primers and protocols used for PCR for cyt-*b* and *dloop*

| **Marker** | **Primers**<br>**(Source)** | **PCR conditions** |
|---|---|---|
| **Cytochrome *b* (1049 bp)** | GluDg: 5′-TGA CTT GAA RAA CCA YCG TTG-3′ (Palumbi 1996)<br>H16460: 5′-CGA YCT TCG GAT TAA CAA GAC CG-3′ (Perdices and Doadrio 2001). | Initial denaturation: (94ºC, 2 min)<br>35 cycles:<br>denaturation (94ºC, 45 s)<br>annealing (53ºC, 45 s)<br>extension (72º, 1.30 min)<br>Final extension: (72ºC, 10 min) |
| **D-loop (599 bp)** | LProF: 5’-AAC TCT CAC CCC TAG CTC CCA AAG-3’ (Meyer et al. 1994)<br>DL623 (5’-GGA ATA GAT ATG TTA TGC ACT TG-3’) (Levin et al. 2003). | Initial denaturation: (95°C, 10 min*)<br>30 cycles:<br>denaturation (95°C, 20 s)<br>annealing (52°C, 30 s)<br>extension (72°C, 1 min)<br>Final extension: (72°C, 5.30 min) |

* Hot Taq Polymerase (Sileks, Moscow).

Table S4. Evolutionary models selected by PartitionFinder for each analyzed gene and for the concatenated data matrix.

| Marker | Total characters | Evolutionary model selected by AIC |
|---|---|---|
| Cyt-*b* (set 1)<br><br>1600000 generations in BI | 1049 | Codon 1st position: SYM+I+G<br><br>Codon 2nd position: HKY<br><br>Codon 3rd position: GTR+G |
| Cyt-*b* + *dloop* (set 2)<br><br>4000000 generations | 1049 Cyt-*b* + 599*dloop* | Codon 1st position: SYM+I+G<br><br>Codon 2nd position: HKY<br><br>Codon 3rd position: GTR+G<br><br>*dloop*: GTR+I+G |

Table S5. Gill raker statistics.

| Forms | Sample size, n | SL, mm | Limits of GR numbers | Coefficient of correlation of GR numbers with SL, mm |
|---|---|---|---|---|
| Generalized | 31 | 73-375 | 14-20 | r = 0.14, NS* |
| Lipped | 8 | 106-459 | 15-18 | r = -0.63, NS |
| Short | 16 | 112-276 | 16-18 | r = 0.15, NS |
| Piscivorous | 14 | 105-426 | 13-16 | r = 0.02, NS |
| *L. jubae* | 15 | 89-349 | 20-26 | r = 0.04, NS |
| Smiling | 13 | 82-360 | 21-24 | r = 0.49, NS |

*NS – not significant.

Table S6. Gut length statistics.

| Forms | Sample size, n | Limits of SL, mm | Variation of gut length as % SL | Coefficient of correlation of GL, % SL, with body length, SL, mm |
|---|---|---|---|---|
| Generalized | 34 | 81-375 | 281-646 | r = 0.72, p<0.01 |
| Lipped | 15 | 106-459 | 281-508 | r = 0.88, p<0.01 |
| Short | 37 | 112-239 | 250-639 | r = 0.31, NS |
| Large-mouthed | 27 | 106-426 | 159-228 | r = -0.23, NS |
| *L. jubae* | 23 | 89-349 | 245-650 | r = 0.70, p<0.01 |
| Smiling | 15 | 82-360 | 195-514 | r = 0.88, p<0.01 |

Table S7. Genetic variation within Genale forms based on *dloop* sequences.

| Form | *n* | *H* | *h*±SD | *π*±SD | *K* | Tajima's *D* | Fu's *Fs* |
|---|---|---|---|---|---|---|---|
| **NEC** | 43 | 31 | 0.98±0.01 | 0.0173±0.0009 | 10.5 | -0.509$^{NS}$ | -11.567* |
| **Large-mouthed** | 33 | 6 | 0.76±0.04 | 0.0036±0.0002 | 2.2 | 1.298$^{NS}$ | 0.796$^{NS}$ |
| **Short** | 20 | 17 | 0.98±0.02 | 0.0182±0.0014 | 11.0 | -0.341$^{NS}$ | -4.657$^{NS}$ |
| **Smiling** | 19 | 6 | 0.74±0.07 | 0.0018±0.0003 | 1.1 | -0.717$^{NS}$ | 0.340$^{NS}$ |
| **Smiling hybrid** | 12 | 12 | 1.00±0.03 | 0.0268±0.0045 | 16.1 | 0.509$^{NS}$ | -1.240$^{NS}$ |
| **L. jubae** | 28 | 25 | 0.99±0.01 | 0.0184±0.0027 | 11.1 | -1.379$^{NS}$ | -9.240* |
| **Lipped** | 26 | 24 | 0.99±0.02 | 0.0182±0.0019 | 11.0 | -1.065$^{NS}$ | -11.763* |

Note: *n* – sample size; *H* – number of haplotypes; *h* – haplotype diversity; *π* – nucleotide diversity (per site); *K* – average number of nucleotide substitutions; SD – standard deviation; $^{NS}$ not significant; * significant at α = 0.02

Table S8. Pairwise $F_{ST}$ values between Genale barb forms based on mtDNA sequence data.

| | **NEC** | **Large-mouthed** | **Smiling** | **Lipped** | ***L. jubae*** | **Short** |
|---|---|---|---|---|---|---|
| **NEC** | | | | | | |
| **Large-mouthed** | 0.59* | | | | | |
| **Smiling** | 0.78* | 0.93* | | | | |
| **Lipped** | 0.01 | 0.57* | 0.76* | | | |
| ***L. jubae*** | 0.62* | 0.76* | 0.39 | 0.60* | | |
| **Short** | 0.01 | 0.61* | 0.77* | 0.01 | 0.62* | |
| **Smiling hybrid** | 0.36 | 0.60* | 0.26 | 0.34 | 0.21 | 0.36 |

*$P$ < 0.05
