## Supplementary figures and images for "Adaptive radiation of barbs of the genus *Labeobarbus* (Cyprinidae) in the East African river"

### Supplementary file 2

#### Fig. S1. Gill raker shapes.

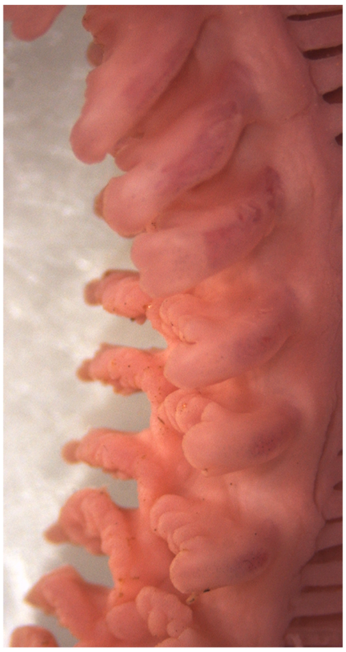

NEC

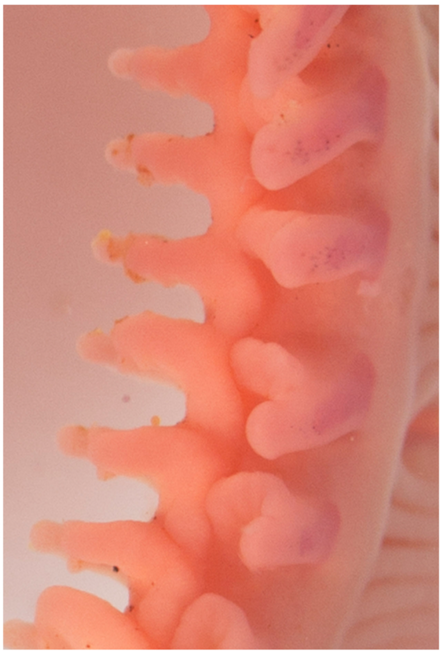

Lipped

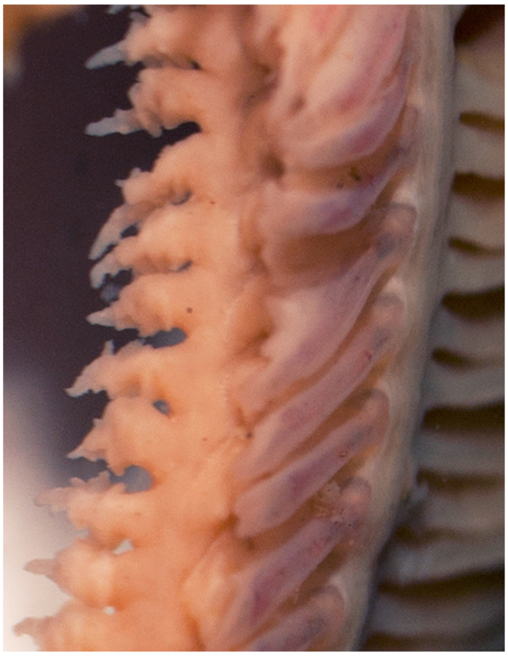

*L. jubae*

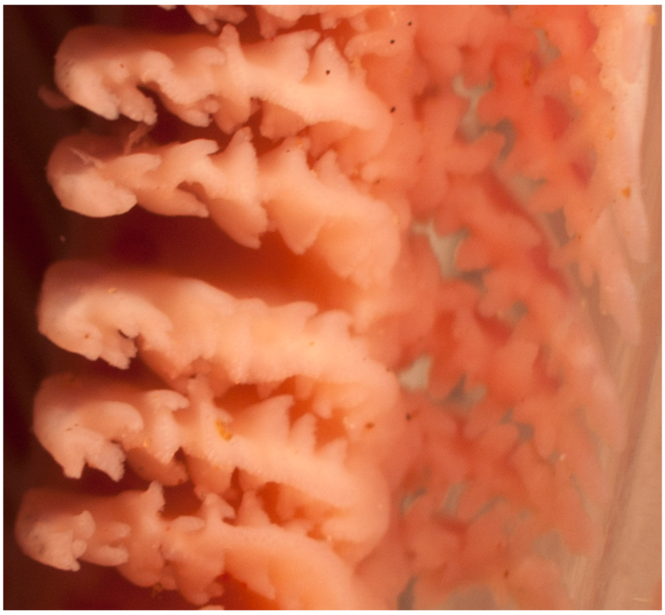

Smiling

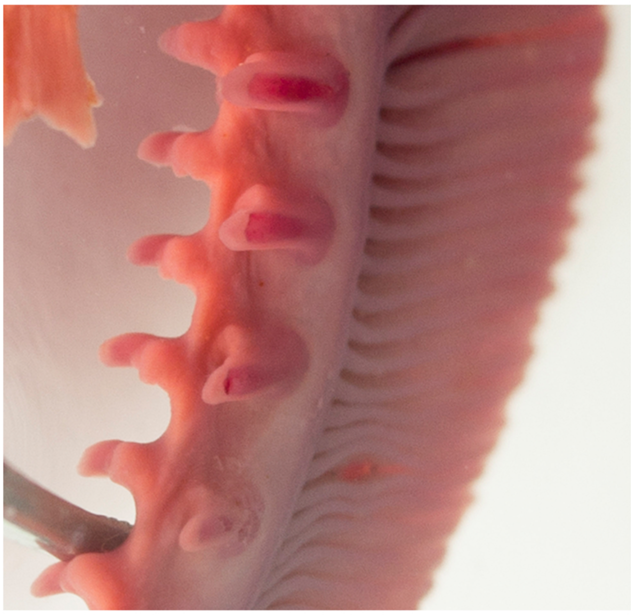

Large-mouthed

### Supplementary file 3

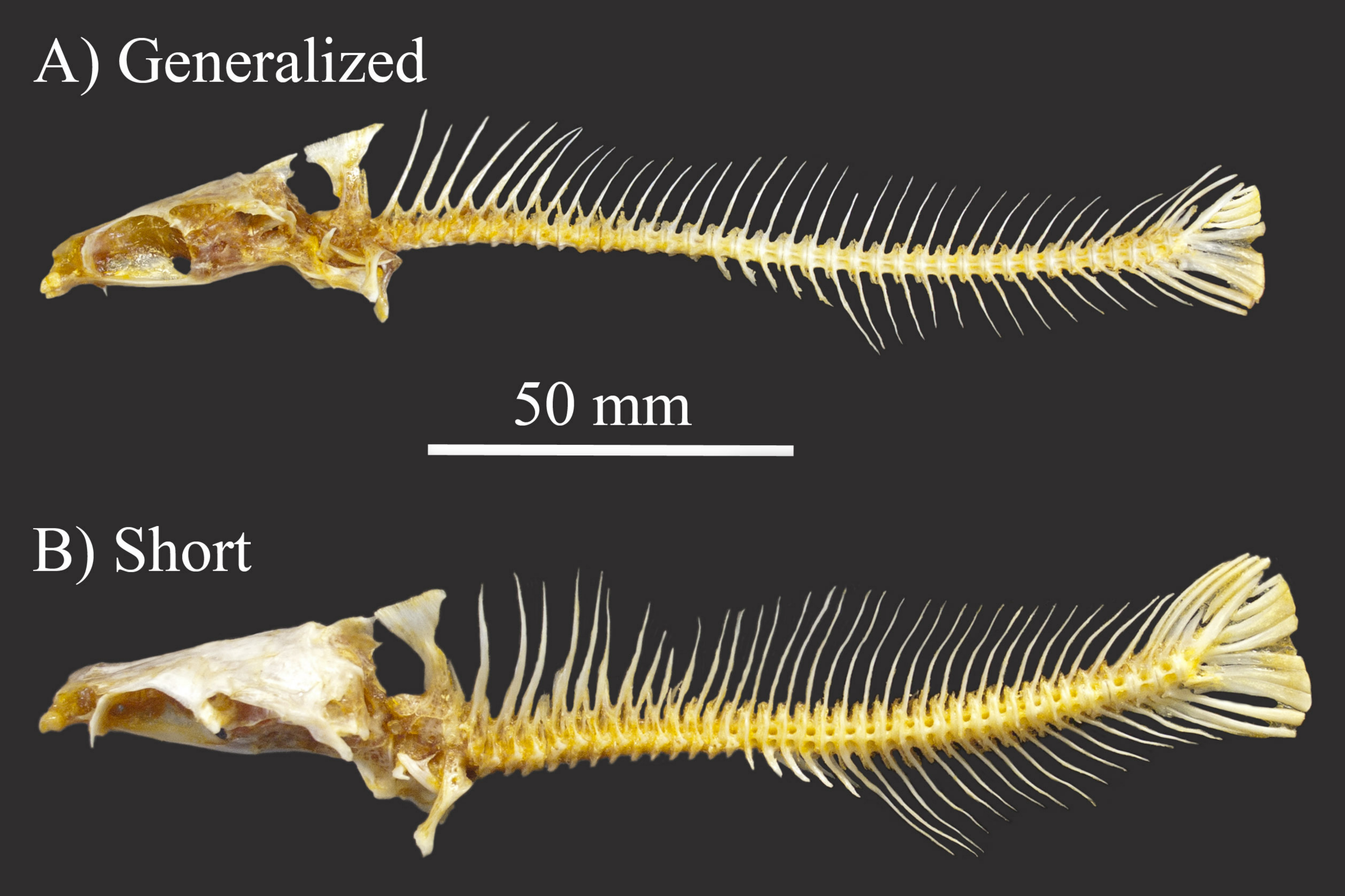

### Supplementary file 4

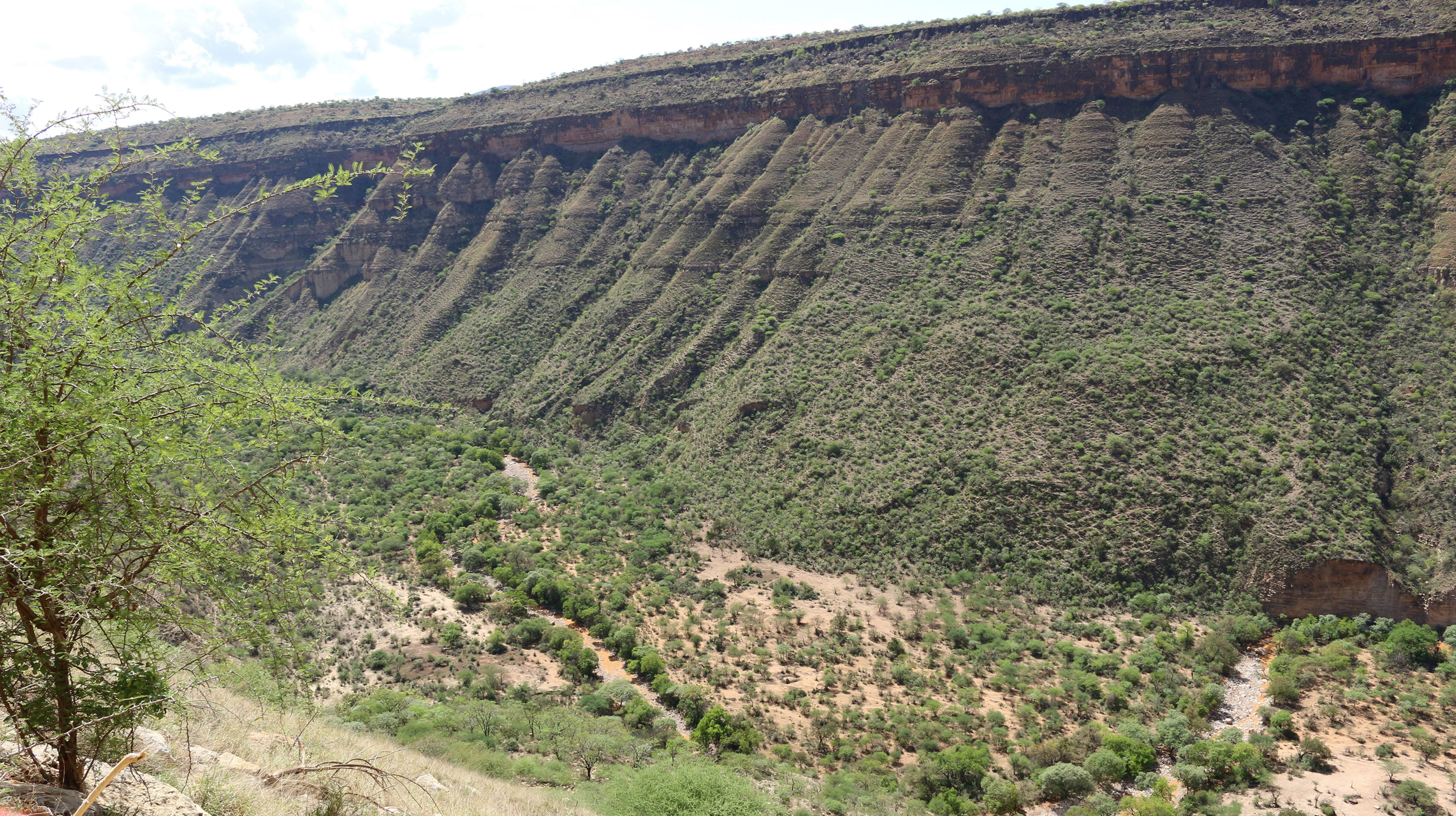

### Supplementary file 5

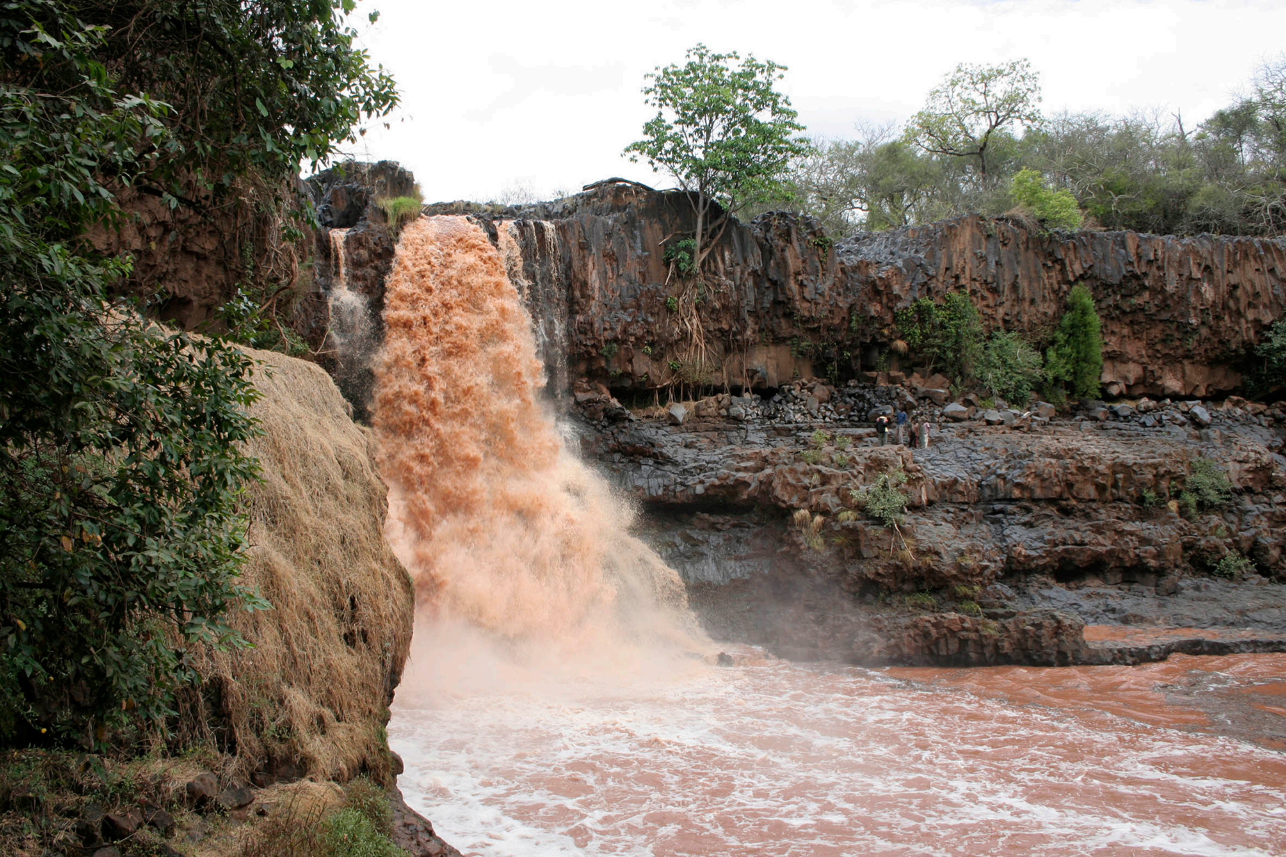
